# Establishing two principal dimensions of cognitive variation in Logopenic Progressive Aphasia

**DOI:** 10.1101/767269

**Authors:** Siddharth Ramanan, Daniel Roquet, Zoë-lee Goldberg, John. R. Hodges, Olivier Piguet, Muireann Irish, Matthew A. Lambon Ralph

## Abstract

Logopenic Progressive Aphasia (LPA) is a neurodegenerative syndrome characterised by sentence repetition and naming difficulties arising from left-lateralised temporoparietal atrophy. Clinical descriptions of LPA largely concentrate on profiling language deficits, however, accumulating evidence points to the presence of cognitive deficits, even on tasks with minimal language demands. Although non-linguistic cognitive deficits in LPA are thought to scale with disease severity, patients at discrete stages of language dysfunction display overlapping cognitive profiles, suggesting individual-level variation in cognitive performance, independent of primary language dysfunction. To address this issue, we used principal component analysis to decompose individual-level variation in cognitive performance in 43 well-characterised LPA patients who underwent multi-domain neuropsychological assessments and structural neuroimaging. The principal component analysis solution revealed the presence of two, statistically independent factors, providing stable and clinically intuitive explanations for the majority of variance in cognitive performance in the syndrome. Factor 1 reflected ‘speech production and verbal memory’ deficits which typify LPA. Systematic variations were also confirmed on a second, orthogonal factor mainly comprising visuospatial and executive processes. Adopting a case-comparison approach, we further demonstrate that pairs of patients with comparable Factor 1 scores, regardless of their severity, diverge considerably on visuo-executive test performance, underscoring the inter-individual variability in cognitive profiles in comparably ‘logopenic’ patients. Whole-brain voxel-based morphometry analyses revealed that speech production and verbal memory factor scores correlated with left middle frontal gyrus, while visuospatial and executive factor scores were associated with grey matter intensity of right-lateralised temporoparietal, middle frontal regions and their underlying white matter connectivity. Importantly, LPA patients with poorer visuospatial and executive factor scores demonstrated greater right-lateralised temporoparietal and frontal atrophy. Our findings demonstrate the inherent variation in cognitive performance at an individual- and group-level in LPA, suggesting the presence of a genuine co-occurring cognitive impairment that is independent of language function and disease severity.

## 1. INTRODUCTION

Logopenic Progressive Aphasia (LPA) is a rare neurodegenerative brain disorder, the canonical features of which centre on language dysfunction, including slowing in spontaneous speech, phonological errors and paraphasias, sentence repetition, sentence comprehension, and word finding difficulties (Gorno-Tempini et al., 2008; Gorno-Tempini et al., 2011; Leyton, Ballard, Piguet, & Hodges, 2014). By contrast, grammatical and articulatory processing and semantic comprehension remain relatively spared in early stages of the disease (Gorno-Tempini et al., 2008). The unique language profile of LPA is proposed to reflect a breakdown in lexical retrieval, phonological working memory and phonological processing, functions that together support sentence repetition, naming, spontaneous speech and working memory (Henry & Gorno-Tempini, 2010; Leyton, Piguet, Savage, Burrell, & Hodges, 2012). Neuroanatomically, the locus of atrophy in early stages of LPA is predominantly left-lateralised and centred on the left inferior parietal lobule, lateral temporal and perisylvian cortical regions surrounding the left superior/middle/inferior temporal gyrus (Gorno-Tempini et al., 2008; Krishnan et al., 2017; Leyton et al., 2012; Rohrer et al., 2010; Teichmann et al., 2013). Over time, however, LPA progresses to affect fronto-insular, medial parietal and temporal cortices, encroaching into right-hemisphere temporoparietal regions (Brambati et al., 2015; Galantucci et al., 2011; Rogalski, Cobia, Harrison, Wieneke, Weintraub, et al., 2011; Rohrer et al., 2013; Tu, Leyton, Hodges, Piguet, & Hornberger, 2016). At a pathological level, the majority of LPA patients (> 90%) present with abnormal levels of cortical β-amyloid, characteristic of Alzheimer’s disease (Chare et al., 2014; Leyton et al., 2011; Rabinovici et al., 2008; Santos-Santos et al., 2018), although recent histopathological and biomarker evidence also points to the presence of non-Alzheimer pathologies in a minority of clinically-diagnosed LPA patients (Bergeron et al., 2018; M. M. Mesulam et al., 2014).

While current classification criteria and clinical descriptions of LPA emphasise the fine-grained characterisation of language dysfunction, mounting evidence points to co-occurring non-linguistic cognitive deficits in this syndrome. Notably, LPA patients have been reported to show impaired processing speed, sustained attention and working memory, and dysexecutive profiles (Butts et al., 2015; Foxe, Irish, Hodges, & Piguet, 2013; Magnin et al., 2013; Rohrer, Rossor, & Warren, 2012). Significant socioemotional dysfunction including loss of empathy and impaired emotion detection abilities has also been documented (Fittipaldi et al., 2019; Hazelton, Irish, Hodges, Piguet, & Kumfor, 2017; Multani et al., 2017). Finally, LPA patients demonstrate significant verbal episodic memory difficulties (Butts et al., 2015; Casaletto et al., 2017; Eikelboom et al., 2018; Win et al., 2017) comparable to that observed in typical Alzheimer’s disease (Ramanan et al., 2016; Ramanan, Marstaller, Hodges, Piguet, & Irish, 2020). While such deficits could manifest simply as a by-product of language and lexical retrieval difficulties in LPA, compromised performance on tasks with minimal language demands suggests otherwise. For example, LPA patients show significant impairments on nonverbal tasks of episodic memory (Ramanan et al., 2016; Ramanan et al., 2020), spatial span (Foxe et al., 2013; Foxe et al., 2016), spatial orientation (Magnin et al., 2013), and visuospatial processing (Butts et al., 2015; Watson et al., 2018), all of which circumvent language demands. Moreover, impairments on nonverbal episodic memory and emotion processing in LPA have been shown to persist when disease severity and language dysfunction are controlled for (Multani et al., 2017; Ramanan et al., 2016). Clinical and carer reports further corroborate these findings, with the majority of LPA patients presenting with visible extra-linguistic general cognitive difficulties (Owens et al., 2018). Further, changes in socio-emotional, attention and memory functions in LPA are detectable 1-3 years prior to spousal recognition of frank expressive language difficulties in patients (Pozzebon, Douglas, & Ames, 2018). Together, these findings argue against language dysfunction as the sole mediator of general cognitive decline in LPA and suggest the presence of genuine co-occurring non-linguistic cognitive deficits.

**Table 1.**
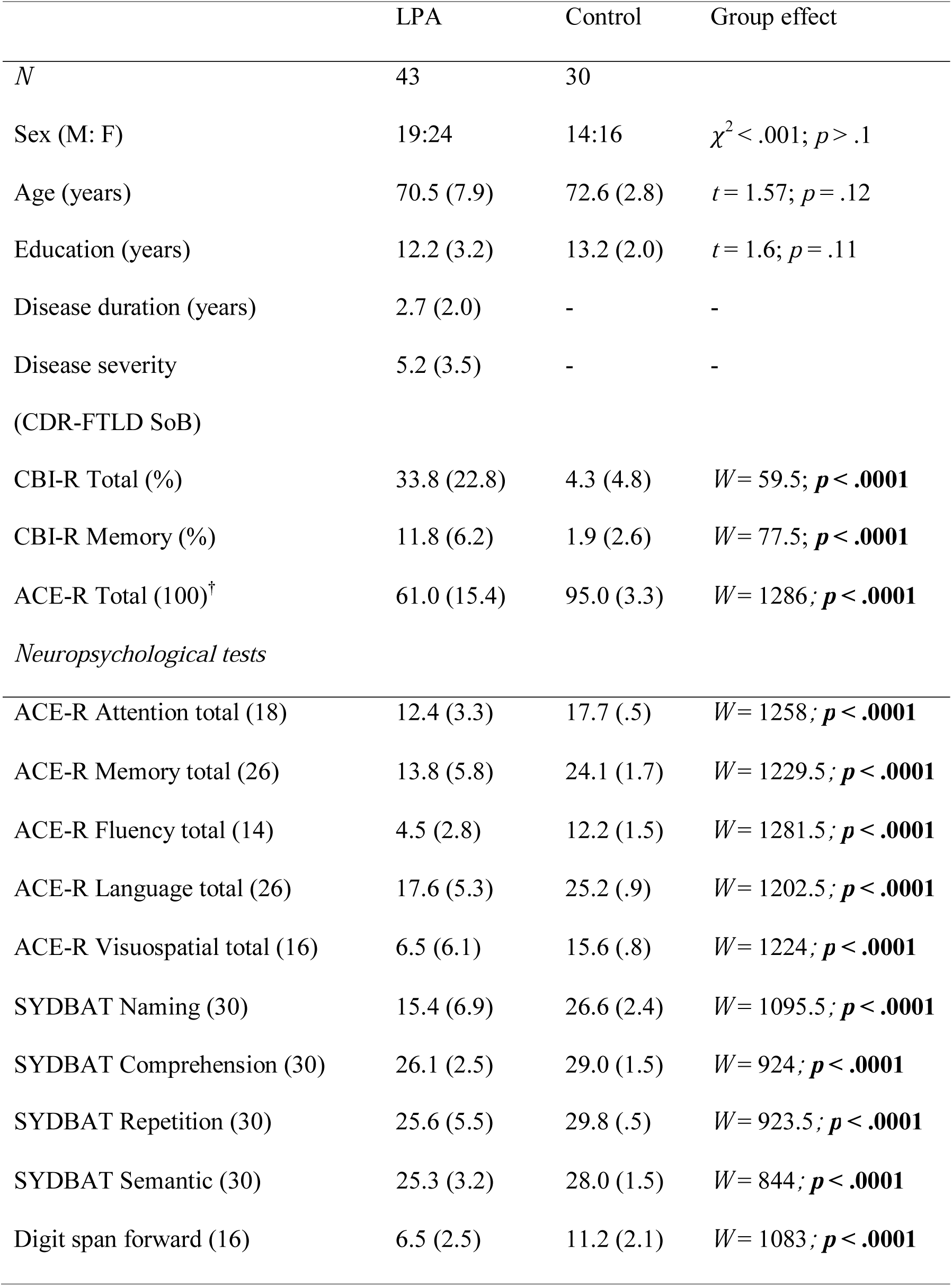
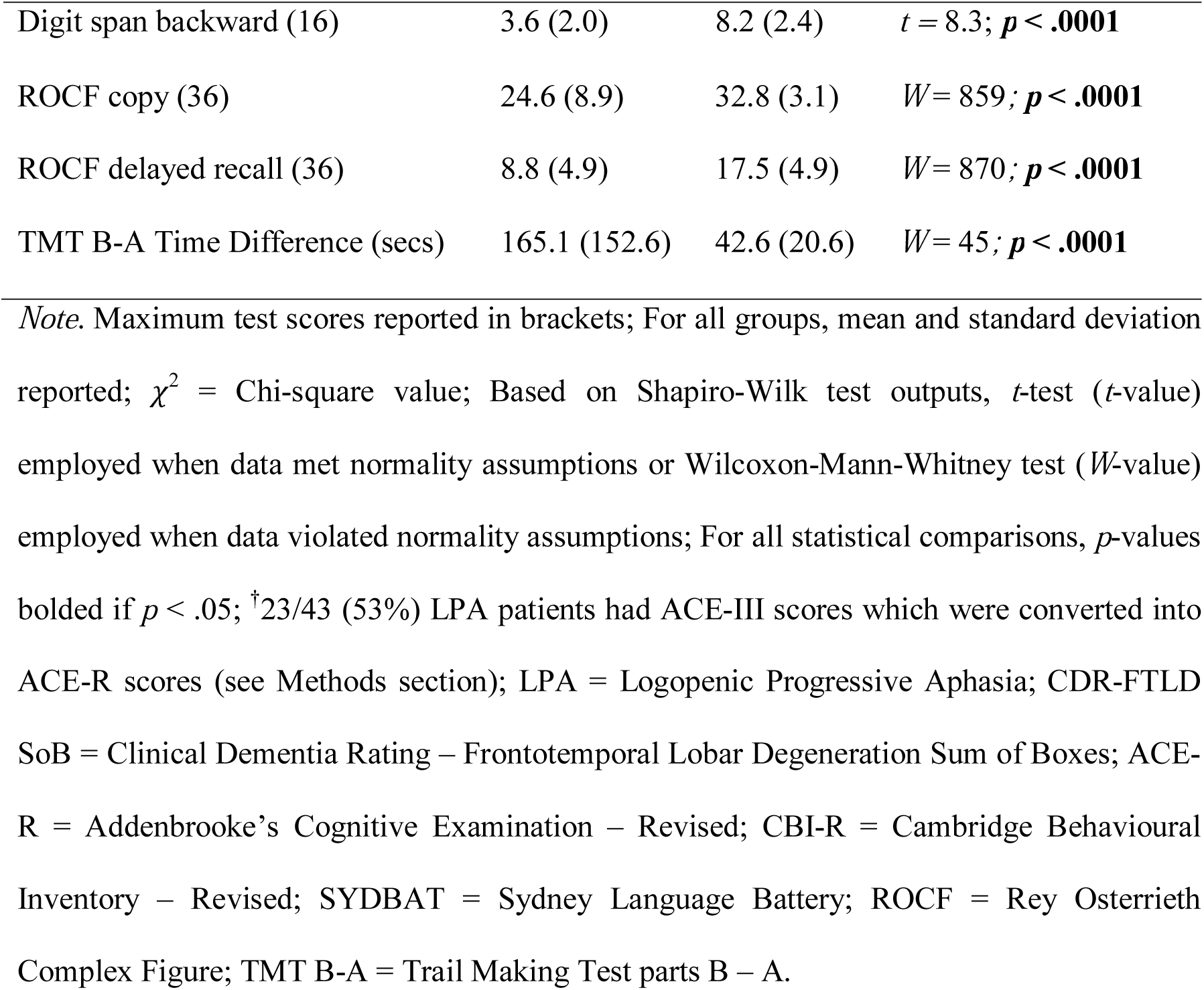
Demographic, clinical, and general neuropsychological assessment performance for all groups.

Given the marked heterogeneity in test performance across cognitive domains and between individual cases in LPA, data-driven approaches hold considerable promise to refine our understanding of this syndrome, as they can simultaneously model systematic variations at a domain- and individual-level. Previous studies in LPA have employed cluster analysis techniques to identify endophenotypes or ‘clusters’ of LPA patients, based on their language performance. These clusters tend to vary primarily along disease severity and degree of aphasia (Leyton et al., 2015; Machulda et al., 2013), and then by level of overall cognitive impairment (Owens et al., 2018). The clinical interpretability of these clusters, however, remains limited for two main reasons. First, endophenotypes of LPA identified purely on the basis of language performance tend to overlap significantly in terms of their overall cognitive performance. This suggests that classifying patients exclusively in terms of language dysfunction masks important variations in general cognitive performance in LPA. Second, when examined relative to other primary progressive aphasia syndromes in the context of language performance, LPA rarely emerges as an independent cluster, instead mingling with other neurodegenerative disorders of language (Hoffman, Sajjadi, Patterson, & Nestor, 2017; Ingram et al., 2019; Maruta, Pereira, Madeira, De Mendonca, & Guerreiro, 2015; Sajjadi, Patterson, Arnold, Watson, & Nestor, 2012). Together, these findings suggest that the current practice of identifying LPA endophenotypes on the basis of language disturbances alone, cannot adequately capture the multidimensional nature of cognitive impairments in this syndrome.

Here, we adopted the hypothesis that the multifaceted cognitive dysfunction in LPA reflects graded variations along multiple, continuous dimensions, rather than strictly defined categorical clusters. Graded approaches have been employed to great effect in the post-stroke aphasia literature, where patients present with variable combinations of expressive and receptive language impairments and co-occurring general cognitive deficits attributable to variations in the size and location of lesions (Butler, Lambon Ralph, & Woollams, 2014; Halai, Woollams, & Lambon Ralph, 2017; Kummerer et al., 2013; Mirman et al., 2015; Ramsey et al., 2017) and more recently in large-scale examinations of frontotemporal lobar degeneration-related syndromes (Murley et al., in press) or variations in semantic dementia/temporal lobe variant of frontotemporal dementia (Ding et al., in press). In particular, principal component analysis (PCA) has been used as a data-driven method to reveal statistically-reliable, graded differences across individual cases, placing them relative to each other within the resultant multidimensional space and, in turn, relating these principal components, rather than individual test scores, to the pattern of the patients’ lesions/atrophy. PCA approaches have been used to ‘compress’ and extract weighted scores from multidimensional data (see e.g., Hoffman et al., 2017; Ramanan et al., 2017), aiding the determination of independence or inter-dependence between cognitive domains. In addition, emergent components from PCA can be used to place participants along a spectrum, enabling characterisation of graded variations between participants across cognitive domains. Accordingly, the emergence of a single, weighted component from the PCA would allude to considerable within-group homogeneity, such that a group varies systematically along only one axis of a multidimensional space. In contrast, the emergence of multiple, statistically orthogonal factors confirm systematic, independent differences in multiple cognitive domains within a patient cohort.

To this end, we employed PCA to explore the neurocognitive architecture of language and general cognitive performance in a large well-characterised sample of LPA patients (*N* = 43). Our primary aims were to reveal the extent of graded variations in cognitive performance within the LPA syndrome, and to use the emergent components to characterise patient performance at the individual-level. We predicted that marked cognitive heterogeneity would be evident, regardless of the severity of language impairments. Finally, we sought to establish the neural substrates of the graded variation in cognitive performance within the LPA syndrome, using voxel-based morphometry.

## 2. MATERIALS AND METHODS

Below, we report how we determined our sample size, all data exclusions, all inclusion/exclusion criteria, whether inclusion/exclusion criteria were established prior to data analyses, all manipulations, and all measures in the study.

### 2.1. Participants

A total of 73 participants were recruited through FRONTIER, the frontotemporal dementia research group at the Brain and Mind Centre, The University of Sydney, Australia. Forty three patients with a clinical diagnosis of LPA, presenting with early anomia, word-finding and sentence repetition difficulties, were included (Gorno-Tempini et al., 2011). Diagnoses were arrived at by consensus among a multidisciplinary team comprising a senior neurologist (J.R.H.), a clinical neuropsychologist, and an occupational therapist, based on comprehensive clinical and neuropsychological examination along with structural neuroimaging. Disease severity for LPA patients was established using the clinician-indexed Frontotemporal Lobar Degeneration-modified Clinical Dementia Rating Sum of Boxes score (CDR-FTLD SoB; Knopman et al., 2008).

Thirty healthy control participants were selected through the research volunteer panel at Neuroscience Research Australia and local community clubs. Controls were matched to patient groups for sex, age, and education, and scored 0 on the CDR-FTLD SoB measure. Healthy controls scored 88 or above on the Addenbrooke’s Cognitive Examination – Revised (ACE-R: Mioshi, Dawson, Mitchell, Arnold, & Hodges, 2006) or its updated counterpart, the Addenbrooke’s Cognitive Examination – III (ACE-III: Hsieh, Schubert, Hoon, Mioshi, & Hodges, 2013) – both of which assess global cognitive functioning. Exclusion criteria for participants included a history of significant head injury, cerebrovascular disease, alcohol and drug abuse, other primary psychiatric, neurological, or mood disorders, and limited English proficiency.

All participants or their Person Responsible provided written informed consent in accordance with the Declaration of Helsinki. This study was approved by the South Eastern Sydney Local Healthy District and The University of New South Wales ethics committees.

### 2.2. General and targeted neuropsychological assessments

Participants underwent extensive neuropsychological testing. Global cognitive functioning was indexed using the ACE-R/ACE-III total score (Hsieh et al., 2013; Mioshi et al., 2006), which includes subtests of attention (max = 18), verbal memory (max = 26), verbal fluency (max = 14), language (max = 26), and visuospatial (max = 16) function. A subset of LPA patients (*N* = 23, ∼53% of the LPA sample) completed the ACE-III (Hsieh et al., 2013). For comparability, their ACE-III subtest scores were transformed to the equivalent ACE-R subtest scores (see So et al., 2018).

Targeted cognitive assessments of language, visuospatial function, memory, and executive functioning were administered. Confrontation naming, single word comprehension, single word repetition, and semantic association were assessed using the Sydney Language Battery (SYDBAT: Savage et al., 2013). Visuo-constructional abilities were assessed using the Copy score (max = 36) of the Rey Osterrieth Complex Figure test (ROCF: Osterrieth, 1944), while the 3-minute delayed recall (max = 36) of the ROCF was used to index nonverbal episodic memory. Auditory attention and working memory were measured using Digit Span Forward and Backward tests, respectively (Strauss, Sherman, & Spreen, 2006). Finally, executive dysfunction was indexed via the Trail Making Test B-A time difference (TMT B-A: Reitan, 1958).

### 2.3. Statistical analyses

Statistical analyses of behavioural data were conducted using a combination of RStudio v3.3.0 (R Core Team, 2016) and MATLAB (The Mathworks Inc., Natick, MA), described below and in Supplementary Methods.

#### 2.3.1. Step 1: Characterising group differences

Group differences on demographic, clinical and neuropsychological performance between LPA and Control groups were explored. For binomially distributed variables (*i.e*., sex), Chi-squared tests were used. For all continuous variables (*i.e*., demographic, clinical, and neuropsychological test measures), normality of distribution was examined using Shapiro-Wilk tests and box-and-whisker plots. Accordingly, *t*-tests or Wilcoxon-Mann-Whitney tests were respectively employed when data met or violated normality assumptions. Two-tailed Pearson’s correlations (*r* values) with false discovery rate correction for multiple comparisons (Benjamini & Hochberg, 1995) were used to examine associations between neuropsychological test performance and clinician-indexed disease severity (CDR-FTLD SoB) in the LPA group. For all analyses of group differences and correlations, an alpha of *p* ≤ .05 was employed.

#### 2.3.2. Step 2: Tabulating and imputing missing data and standardizing scores

All subsequent statistical analyses were conducted in the LPA group. As PCA algorithms operate on standardized datasets with no missing variables, the frequency of missing neuropsychological data was first tabulated and plotted for subsequent imputation (Supplementary Figure 1). Across all neuropsychological test measures, the LPA group had a total of 4.8% missing data with the majority of patients (17/43 LPA, *i.e*., 39.5% of LPA group) missing TMT B-A data (Supplementary Figure 1). All available data were converted into percentages (detailed in Supplementary Methods) and this final dataset was used for imputation.

Missing data were imputed using a probabilistic PCA using *k*-fold cross-validation approach (with *k* = 4; detailed in Supplementary Methods). Briefly, this approach offers improved stability as compared to list-wise exclusion of rows with missing data, while simultaneously guarding against overfitting of imputed data points (unlike imputation of group mean) (see Ilin & Raiko, 2010; Tipping & Bishop, 1999). The output was a ‘full’ dataset with no missing values.

#### 2.3.3. Step 3: Identifying principal cognitive factors

The final ‘full’ standardized dataset was entered into an orthogonally rotated (varimax) PCA. Varimax rotation facilitates interpretations of PCA output by maximising the dispersion of factor loadings between components, allowing for little variance to be shared commonly between emergent components. In line with standard approaches (Jolliffe, 2002), factors with an eigenvalue of 1.0 and above were extracted. Each factor was given a label reflecting the majority of tests loading heavily (*i.e*., loadings > 0.5) on that factor.

It must be noted that factor names are simply shorthand labels that reflect the majority of cognitive tests loading onto that particular factor, and by no means reflect the entirety of cognitive processes that underpin performance on each test loading onto that particular factor. Individual patient scores on each factor were extracted and used as orthogonal covariates in subsequent neuroimaging analyses. In addition, we projected the lower bound of normality (*i.e*., −1.96 standard error of the mean) from the control data into the patients’ PCA space to facilitate behavioural interpretation of patient factor scores relative to control test performance (detailed in Supplementary Methods). Finally, associations between disease severity, disease duration and emergent factor scores were examined using two-tailed Pearson’s correlations.

#### 2.3.4. Step 4: Computing deviations from expected cognitive performance

As PCA results are one-step removed from raw test scores, we used PCA factor scores to predict each patient’s ‘ideal’ test performance and compared their predicted and raw test neuropsychological performance (adopting the approach used in Lambon Ralph, Patterson, Graham, Dawson, & Hodges, 2003). This approach translates information from the PCA space back into readily-comprehensible predicted test scores, allowing for direct and intuitive comparisons of expected and actual test performance between LPA patients.

Our PCA generated two orthogonal factors. Tests that loaded heavily on Factor 1 resembled measures on which LPA patients typically show early deficits (*e.g*., naming, repetition, verbal working memory and short-term memory). By contrast, tests that loaded heavily on the orthogonal factor (Factor 2) reflected measures on which performance is traditionally thought to be affected in later stages of LPA (*e.g*., visuospatial, executive and comprehension measures). We therefore treated each patient’s Factor 1 score as a simple metric of how ‘logopenic’ they are and used these scores to predict test performance on neuropsychological measures loading differentially on Factors 1 and Factors 2. This comparison would demonstrate how comparably logopenic patients (with similar Factor 1 scores) diverge on test measures posited to be relatively preserved, until later stages of LPA.

To do this, we first visually identified and selected four pairs of LPA patients (denoted using pairwise matching colours in Figure 1). Each pair was carefully selected so that they i) had comparable scores on Factor 1 but, ii) diverged on Factor 2 scores, and iii) were sampled across varying Factor 1 scores to reflect the spread of distribution along the *x*-axis (see Lambon Ralph et al., 2003 for similar analyses). Following pair selection, we employed a series of linear regression analyses using Factor 1 scores to predict performance on select neuropsychological tasks that loaded heavily on Factor 1 (SYDBAT Naming and Repetition, and Digit Span Forward) and Factor 2 (SYDBAT Comprehension and ROCF Copy and Delayed Recall). Each pair’s predicted scores were then visually compared to their raw neuropsychological test scores.

**Fig 1.**
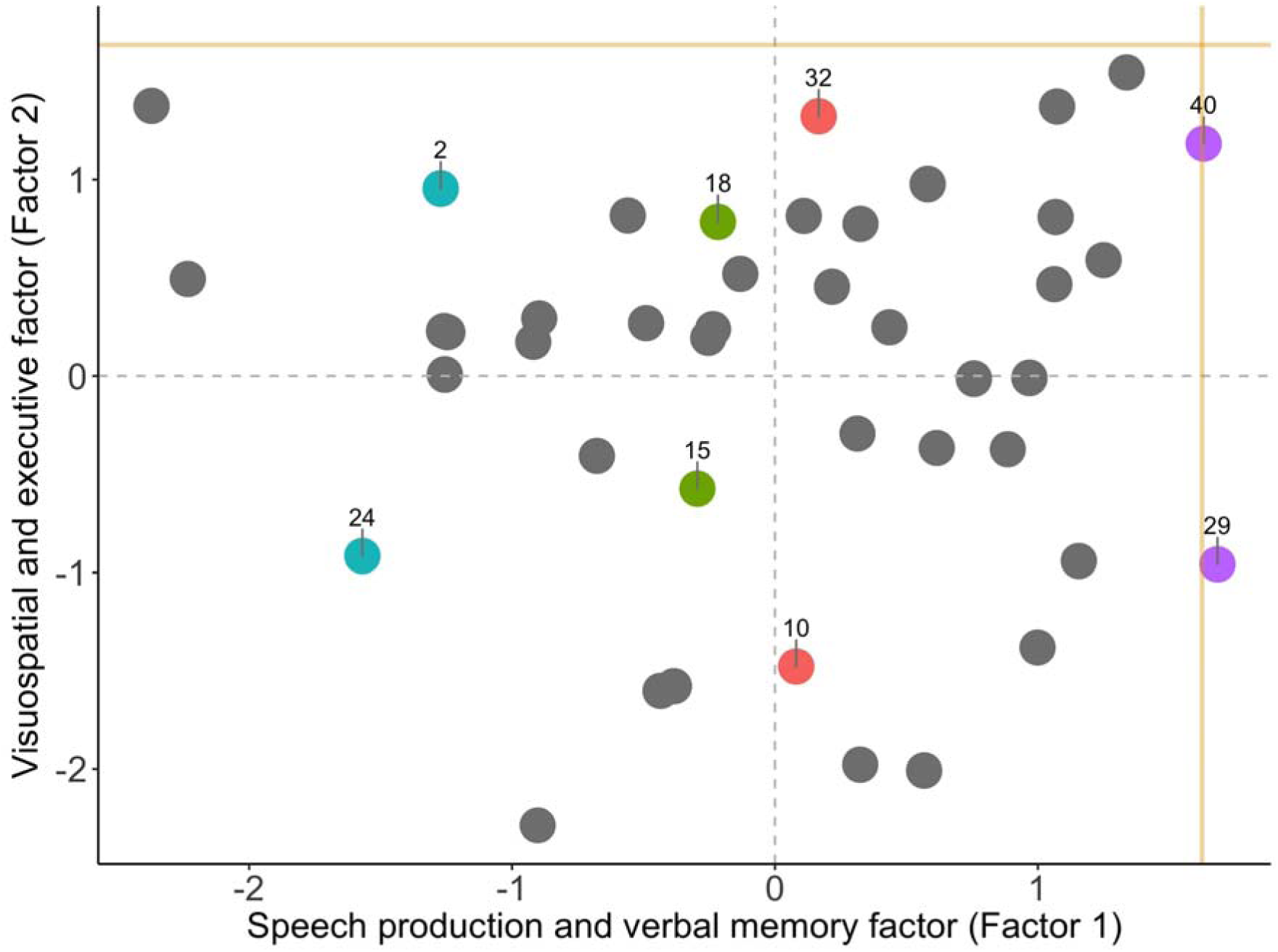
Factor scores of LPA patients on the speech production and verbal memory factor (*i.e*., Factor 1) and visuospatial and executive factor (*i.e*., Factor 2) emerging from the varimax-rotated Principal Component Analysis. Coloured data points indicate individual patients who were examined in pairwise fashion in subsequent statistical analyses, with matching colours denoting patient pairs of interest. Gold lines indicate lower bound of normality (−1.96 standard error from the mean) as estimated from the Control group (calculation detailed in Supplementary Methods). LPA = Logopenic Progressive Aphasia.

### 2.4. Image acquisition

Sixty-three participants (35 LPA and 28 Controls) underwent structural T_1_-weighted brain MRI using a 3T Philips MRI scanner with standard quadrature head coil (eight channels). All 3D T_1_-weighted images were acquired using the following sequences: coronal acquisition, matrix 256 × 256 mm, 200 slices, 1mm isotropic voxel resolution, echo time/repetition = 2.6/5.8 msec, flip angle α=8°.

We used combined grey and white matter voxel-based morphometry (VBM) to account for co-occurring cortical grey and subcortical white matter changes that are prototypical of neurodegenerative disease syndromes such as LPA (Brambati et al., 2015). Such a method has been employed in populations presenting with diffuse, co-occurring grey and white matter changes such as healthy ageing (Giorgio et al., 2010), post-stroke aphasia (Halai et al., 2017), and frontotemporal lobar degeneration syndromes (Lansdall et al., 2017; Murley et al., in press). VBM analyses were conducted using Statistical Parametric Mapping software (SPM12: Wellcome Trust Centre for Neuroimaging, https://www.fil.ion.ucl.ac.uk/spm/software/spm12/). Full details of the standard pre-processing pipeline are provided in Supplementary Methods.

### 2.5. VBM analyses

#### 2.5.1. Whole-brain changes in grey and white matter intensity

Voxelwise differences of grey and white matter intensity between LPA and Control groups were assessed using independent *t*-tests, with age and total intracranial volume included as nuisance variables. Clusters were extracted, corrected for Family-Wise Error at *p* < .01 with a cluster threshold of 100 contiguous voxels. Emergent clusters were subsequently binarized into a mask that was used to compute voxel-level variance in grey and white matter intensity (see below).

#### 2.5.2. Variance in grey and white matter intensity across participants

VBM correlation analyses are entirely constrained by variations in voxel-level intensity and test performance. In the context of progressive diseases, this means that highly atrophic regions that subsequently have uniformly low voxel-level variance are unlikely to emerge in the correlation analyses as they are consistently affected across cases. These regions, nevertheless, could be critical to explaining the observed behavioural profile and therefore, it is important to interpret VBM results in the context of whole-brain voxel-level variance. To complement our atrophy analyses, we therefore computed voxel-level inter-subject variance maps of grey and white matter intensity for all participants. The resultant whole-brain images were further masked to consider only clusters emerging in our atrophy analyses. As before, age and total intracranial volume were regressed out as nuisance variables prior to extracting variance maps.

#### 2.5.3. Grey and white matter intensity changes in patients stratified on factor scores

We further investigated whole-brain changes in grey and white matter intensity in patients with ‘low’ and ‘high’ factor scores. Patients were stratified into two folds on either end of a zero score on Factor 1 and Factor 2 each (see Supplementary Figure 2). Stratifying on Factor 1 resulted in 15 patients with negative (low) and 20 patients with positive (high) scores while stratifying on Factor 2 resulted in 16 patients with negative (low) and 19 patients with positive (high) scores (Supplementary Figure 2). Patients split on Factor 1 scores had comparable Factor 2 scores and vice versa (both *p* values > .1). When compared to patients with higher Factor 1 scores, those with lower Factor 1 scores had greater disease severity (*t* = 2.52; *p* = .016), whilst the difference of disease duration was borderline (*t* = 1.9; *p* = .065). In contrast, no significant group differences were noted on disease severity (*t* = .37; *p* = .70) and disease duration (*t* = −1.19; *p* = .24) between patients split on Factor 2 scores. Regression models with separate directional contrasts (*i.e*., independent *t*-tests) were used to assess differences in cortical grey matter and subcortical white matter intensities between LPA subgroups (*i.e*., high and low scorers) on each Factor score, with age and total intracranial volume included as nuisance variables. Clusters were extracted at *p* < .001, uncorrected, with a cluster threshold of 100 contiguous voxels.

#### 2.5.4. Correlations with PCA-generated factor scores

Finally, correlation analyses within the LPA group (*N* = 35) were employed to examine associations between whole-brain grey and white matter intensity and PCA-generated factor scores. A correlation-only statistical model was implemented for additional statistical power, using *t*-contrasts to measure associations between grey and white matter intensity and PCA-generated factor scores. Age and total intracranial volume were included as nuisance covariates in the analyses. Anatomical locations of statistical significance were overlaid on the MNI standard brain with maximum co-ordinates provided in MNI stereotaxic space. Clusters were extracted using a threshold of *p* < .001 uncorrected for multiple comparisons with a cluster threshold of 100 contiguous voxels.

### 2.5. Data availability

The ethical requirement to ensure patient confidentiality precludes public archiving of our data. Researchers who would like to access the raw data should contact the corresponding authors, who will liaise with the ethics committee that approved the study, and accordingly, as much data that are required to reproduce the results will be released to the individual researcher. The code used for this project has been made available for review on the Open Science Framework website (https://osf.io/bn534/). No part of the study procedures or analyses were preregistered prior to the research being undertaken.

## 3. RESULTS

### 3.1. Demographic, clinical, and neuropsychological test performance

Demographic, clinical and neuropsychological scores are presented in Table 1. No significant group differences emerged for sex distribution, age, and education (all *p* values > .1). LPA patients performed significantly worse than Controls on measures of global cognition, as well as targeted neuropsychological assessments of episodic memory, semantic naming and comprehension, single-word repetition, visuo-constructional abilities and executive function (all *p* values < .0001; see Table 1). Carers of LPA patients reported significant changes in behaviour and memory on the CBI-R relative to Controls (both *p* values < .0001). These profiles are in keeping with previous descriptions of the LPA cognitive profile (Butts et al., 2015; Magnin et al., 2013; Ramanan et al., 2016; Watson et al., 2018).

### 3.2. Correlations between disease severity and neuropsychological test performance

LPA Digit Span Forward performance correlated with disease severity scores on the CDR-FTLD SoB (*r* = -.39; *p* = .010). No other significant correlations emerged between neuropsychological test performance and disease severity in LPA (all *p* values ≥ .059; see Supplementary Table 1).

### 3.3. Identifying principal cognitive factors

Factors and individual test loadings from the varimax-rotated PCA output are displayed in Table 2, while factor loadings for all LPA patients are displayed in Figure 1 and Supplementary Table 2. The sample size was considered adequate for the analysis (Kaiser-Meyer-Olkin statistic = 0.63). The PCA solution revealed two independent, orthogonal factors that together accounted for 56.4% of the total variance (Factor 1 = 41.8% and Factor 2 = 14.6% of total variance) in LPA cognitive performance. The extraction of a three or four component solution, by contrast, aided little additional explanatory power (Factor 3 = 9.4% and Factor 4 = 7.6%) and only served to split the measures loading on Factor 2 into further independent principal components. We, therefore, chose the two-factor solution for its stability, explanatory power, and clinical intuitiveness in explaining LPA cognitive performance.

**Table 2.** Factor loadings for neuropsychological test measures on the omnibus varimax-rotated Principal Component Analysis.

| Neuropsychological tests | **Factor 1** | **Factor 2** |
| --- | --- | --- |
|  | Speech production and verbal memory factor | Visuospatial and executive factor |
| ACE-R Language Total | **.849** | .114 |
| Digit Span Forward | **.801** | .053 |
| SYDBAT Repetition | **.788** | .036 |
| SYDBAT Naming | **.687** | .200 |
| ACE-R Memory Total | **.662** | .290 |
| Digit Span Backward | **.604** | .405 |
| ROCF Copy | .111 | **.918** |
| SYDBAT Semantic Association | .196 | **.801** |
| SYDBAT Comprehension | .056 | **.782** |
| TMT B-A Time Difference | .146 | **.727** |
| ROCF Delayed Recall | .380 | **.660** |
| ACE-R Attention Total | .444 | **.582** |
| ACE-R Visuospatial Total | .437 | .322 |
| ACE-R Fluency Total | .364 | .283 |

*Note.* Tests that load heavily (loadings > .5) on each factor are indicated in bold. Scores for only LPA patients were entered into the Principal Component Analysis; LPA = Logopenic Progressive Aphasia; ACE-R = Addenbrooke's Cognitive Examination – Revised; SYDBAT = Sydney Language Battery; ROCF = Rey Osterrieth Complex Figure; TMT B-A = Trail Making Test parts B-A.

Factor 1 loaded heavily on tests of verbal memory (ACE-R Memory Total), phonological working memory (Digit Span Forward and Backward, SYDBAT Repetition), naming (ACE-R Language Total, SYDBAT Naming), and repetition (SYDBAT Repetition and Digit Span Forward and Backward) (Table 2). Together, these tests index cognitive processes that are canonically impaired in LPA; therefore, we labelled this factor the ‘speech production and verbal memory factor’.

Our PCA analyses further suggested the presence of an orthogonal set of variations on a second factor. Factor 2 mainly loaded on measures of executive (Trails Time Difference), attention (ACE-R Attention Total), and visuospatial (ROCF Copy and Delayed Recall) abilities. In addition, the SYDBAT Comprehension subtest performance also loaded onto this factor. For brevity, we refer to this factor as the ‘visuospatial and executive factor’. Importantly, patients with both high and low Factor 1 scores exhibited uniform variation on Factor 2 scores and this variation was noted both proximally and distally from the lower bound of normal control performance (Figure 1). Together, these findings suggest that Factor 2 is not solely accounted by the emergence of additional impairments with disease severity, but instead reflects systematic variations on visuospatial and executive performance in LPA patients.

In summary, our PCA pointed to the existence of two orthogonal sets of variations in neuropsychological performance in LPA. While the first factor resembles the classic language profile of LPA, the uniform distribution of scores on Factor 2 suggests a co-occurring primary disruption of visuospatial and executive processes in this syndrome.

### 3.4. Associations between factor scores, disease severity, and disease duration

No significant correlations were found between disease severity (CDR-FTLD SoB) and scores on the speech production and verbal memory factor (Factor 1; *r* = -.25; *p* = .1) or visuospatial and executive factor (Factor 2; *r* = -.16; *p* > .1) (Supplementary Figure 3). In contrast, there was a significant correlation between disease duration and the speech production and verbal memory factor (Factor 1; *r* = -.53; *p* = .0002), but not with the visuospatial and executive factor (Factor 2; *r* = .13; *p* > .1) (Supplementary Figure 4). The lack of strong and statistically significant associations, especially on Factor 2, supports our PCA findings of systematic variations on visuospatial and executive test performance, regardless of the disease severity or disease duration of LPA patients.

### 3.5. Comparably logopenic cases diverge on visuospatial and executive performance

In a second step, we aimed to demonstrate how patients who present as ‘comparably logopenic’ can show divergent visuospatial and executive performance. For this, we first chose LPA patient pairs with comparable Factor 1 scores (*i.e*., coloured pairs in Figure 1). We used their Factor 1 scores to predict neuropsychological performance on selected measures loading differentially on Factor 1 and 2. These predicted scores were then compared to their actual raw neuropsychological performance (Figures 2 and 3).

**Fig 2.**
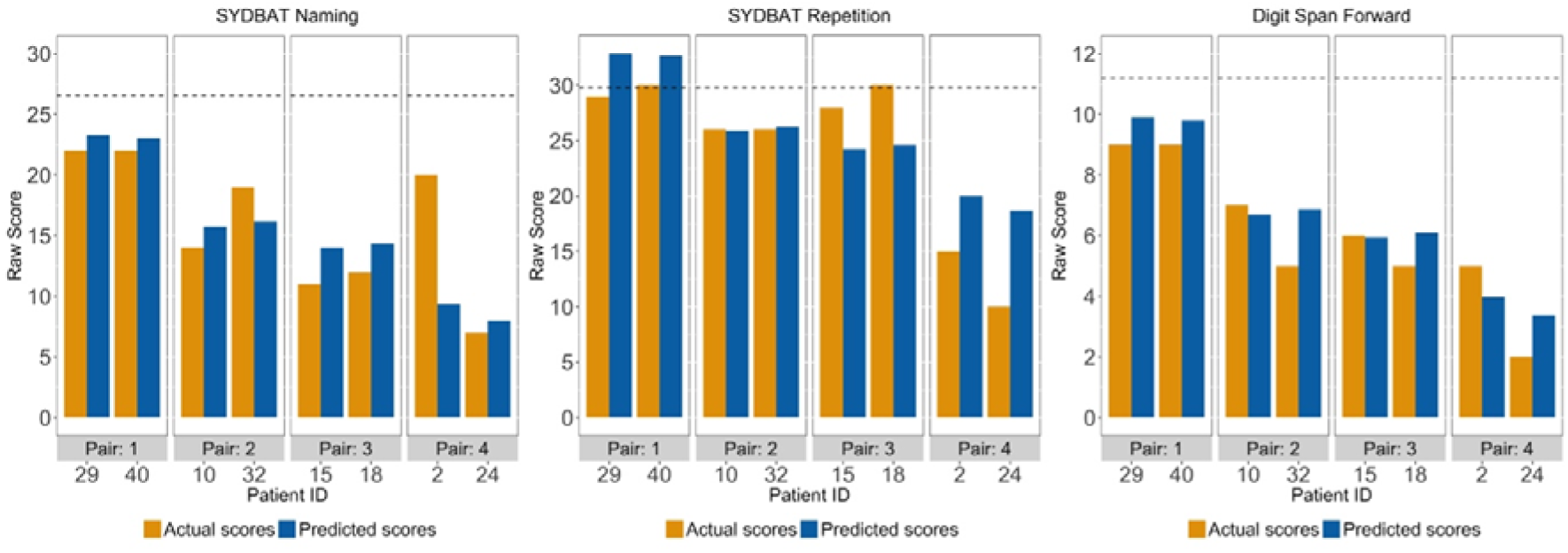
Predicted and actual scores for LPA patient pairs on three example tests loading on the speech production and verbal memory factor (*i.e*., Factor 1) from the varimax-rotated Principal Component Analysis. Dotted lines for each test indicate actual Control mean. SYDBAT = Sydney Language Battery; LPA = Logopenic Progressive Aphasia.

**Fig 3.**
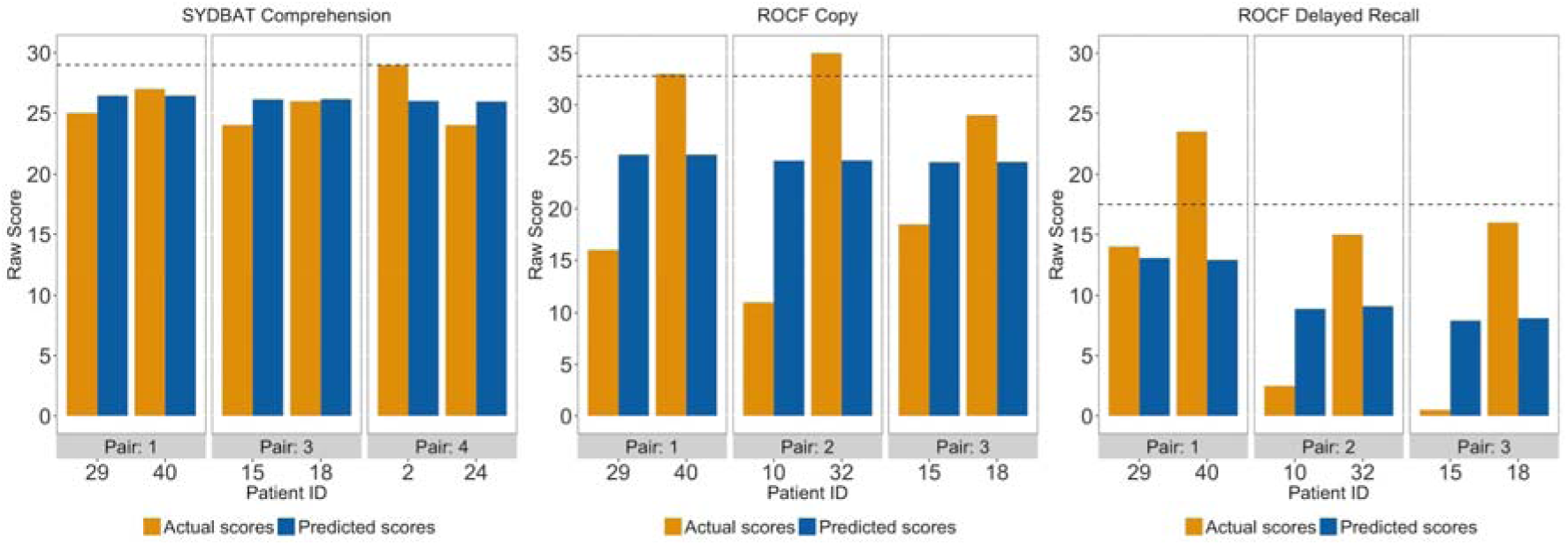
Predicted and actual scores for LPA patient pairs on three example tests loading on visuospatial and executive factor (*i.e*., Factor 2) from the varimax-rotated Principal Component Analysis. Only three pairs presented as one patient from one of the excluded pairs was missing data on the SYDBAT Comprehension or the ROCF measures. Dotted lines for each test indicate actual Control mean. SYDBAT = Sydney Language Battery; ROCF = Rey Osterrieth Complex Figure; LPA = Logopenic Progressive Aphasia.

For tests loading on the speech production and verbal memory factor (Factor 1), predicted and actual scores were nearly similar across all patient pairs (except for pair 4 on SYDBAT Naming) (Figure 2). This pattern confirmed our prediction as comparably ‘logopenic’ patients should display near identical performance on cognitive tasks that are prototypically affected in the LPA syndrome. By contrast, patients displaying comparable ‘logopenic’ presentations (on Factor 1) diverged considerably in terms of predicted and actual scores on visuo-executive measures (Factor 2: ROCF Copy and Delayed Recall) (Figure 3). At an individual level, these findings support the view that while two LPA patients can manifest with comparable severity of ‘logopenic’ symptoms, considerable heterogeneity exists in terms of co-occurring visuospatial and executive impairment in this syndrome.

### 3.6. VBM results

#### 3.6.1. Group differences in grey and white matter intensity

Group differences in grey and white matter intensity are presented in Supplementary Table 3 and Figure 4A. Relative to Controls, the LPA group displayed significant reductions in grey and white matter intensity predominantly in temporo-parietal regions including bilateral superior/middle/inferior temporal gyri (left > right) and bilateral angular and supramarginal gyri (left > right) and underlying white matter bundles, namely the inferior longitudinal and inferior fronto-occipital fasciculi. This cluster extended medially through the underlying white matter into posterior/middle cingulate cortices (left > right) and subcortically into bilateral hippocampi (across the longitudinal axis) and parahippocampal gyri through the cingulum bundle, further into the bilateral thalami, amygdalae (all left > right) and the underlying anterior thalamic radiation (Figure 4A). Relative to Controls, the LPA group further demonstrated reduced grey and white matter intensity in frontal regions such as bilateral insular and superior/middle frontal cortices (both left > right) and underlying white matter connections from the superior longitudinal fasciculus, extending to the right orbitofrontal cortex and its underlying white matter connections into the bilateral temporal poles through the uncinate fasciculus (Figure 4A). These patterns of atrophy are in line with previous descriptions of cortical grey matter and subcortical white matter damage in LPA (Galantucci et al., 2011; Gorno-Tempini et al., 2004; Rogalski et al., 2014; Rohrer et al., 2013; Tu et al., 2016).

**Fig 4.**
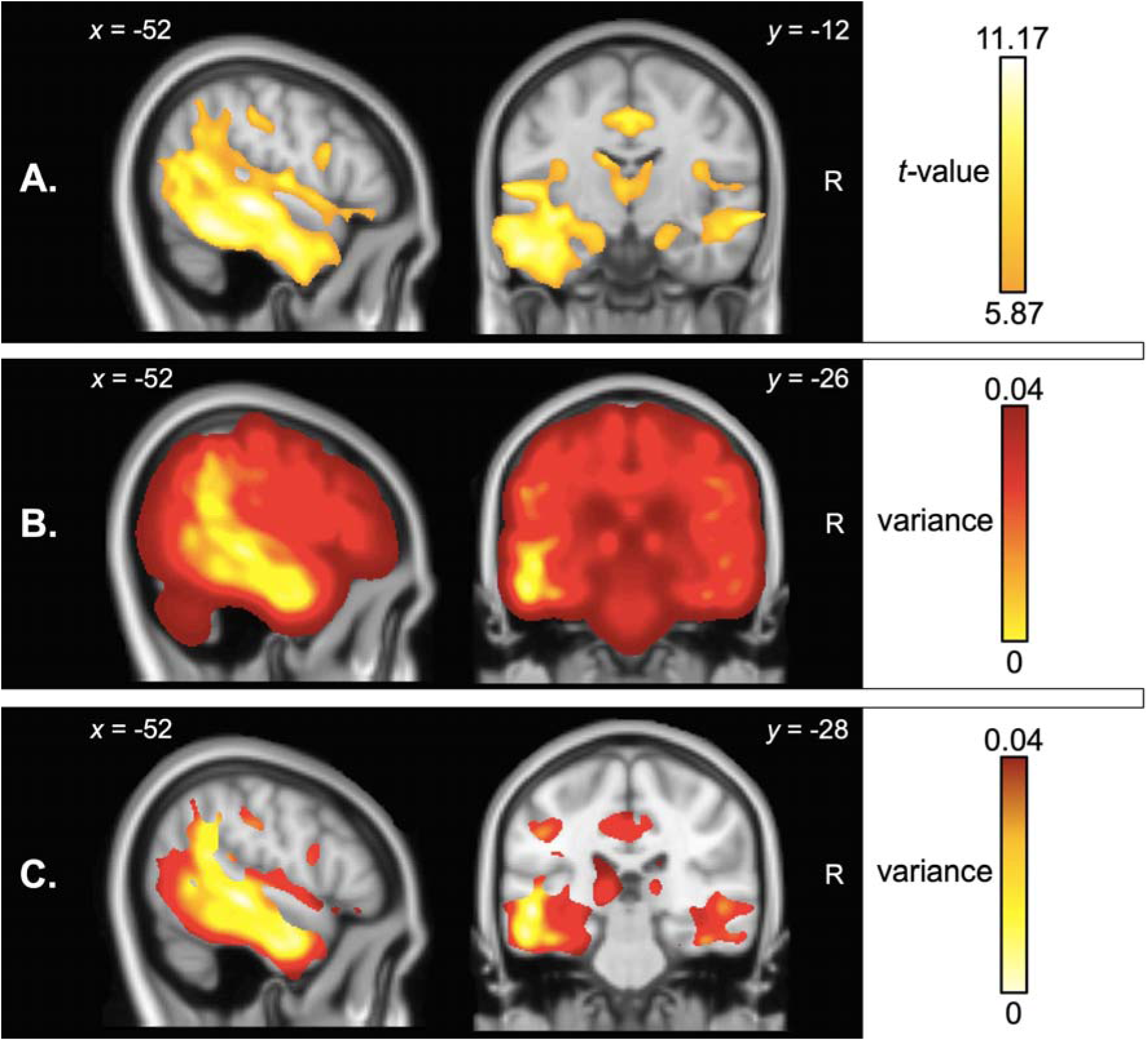
Voxel-based morphometry analyses showing A) regions of significant grey and white matter intensity reduction in LPA compared to Controls, B) voxel-wise variance in grey and white matter intensity in LPA compared to Controls, and C) voxel-wise variance in regions of peak atrophy (computed within a mask of regions emerging from the atrophy analysis in panel A). Coloured voxels in panel A indicate regions that emerged significant in the voxel-based morphometry analyses at *p* < .01 corrected for Family-Wise Error with a cluster threshold of 100 contiguous voxels. Age and total intracranial volume were included as covariates in all analyses. Clusters are overlaid on the Montreal Neurological Institute (MNI) standard brain with *x* and *y* co-ordinates reported in MNI standard space. R = Right; LPA = Logopenic Progressive Aphasia.

#### 3.6.1. Mapping voxel-wise variance in grey and white matter intensity

Visual inspection of variance maps revealed that variance in whole-brain grey and white matter intensity was lowest in left peri-sylvian regions, typically affected in the earliest stages of LPA (Figure 4B). Examining variance within regions of peak atrophy revealed that the area of lowest variance was centred on the left superior/middle temporal gyrus extending into the left temporoparietal junction and inferior parietal cortex; regions that together demonstrated maximal atrophy (*i.e*., lowest grey and white matter intensity) in LPA (Figure 4C). By contrast, regions located at the ‘edges’ of the atrophy clusters and beyond demonstrated maximal variance.

#### 3.6.3. Grey and white matter intensity changes in patients stratified on factor scores

Group differences in grey and white matter intensity are presented in Supplementary Table 4 and Supplementary Figure 5. Direct comparison of LPA subgroups revealed that compared to cases with higher visuospatial and executive factor scores (Factor 2), patients with lower visuospatial and executive factor scores demonstrated greater grey and white matter intensity reduction in predominantly right temporoparietal regions including angular gyrus and supramarginal gyri connecting to superior/middle temporal gyri through the subcortical component of the middle/inferior longitudinal fasciculus. This cluster extended medially towards the right precuneus, posterior cingulate, occipital cortices. This cluster further extended rostrally towards right frontal regions such as middle/inferior frontal gyrus and middle cingulate gyrus through the subcortical cingulum bundle and superior longitudinal fasciculus tract, and subcortically towards the right parahippocampal regions and fusiform gyrus. Additionally, a relatively smaller cluster centred around the left angular gyrus, precuneus, and underlying superior/inferior longitudinal fasciculus bundles was noted. No significant results emerged for the reverse contrast or for contrasts comparing high and low scores on the speech production and verbal memory factor (Factor 1) (Supplementary Table 4).

#### 3.6.4. Neural correlates of principal cognitive factors

Associations between grey and white matter intensity and factor scores in the LPA group are displayed in Figure 5 and Table 3.

**Table 3.**
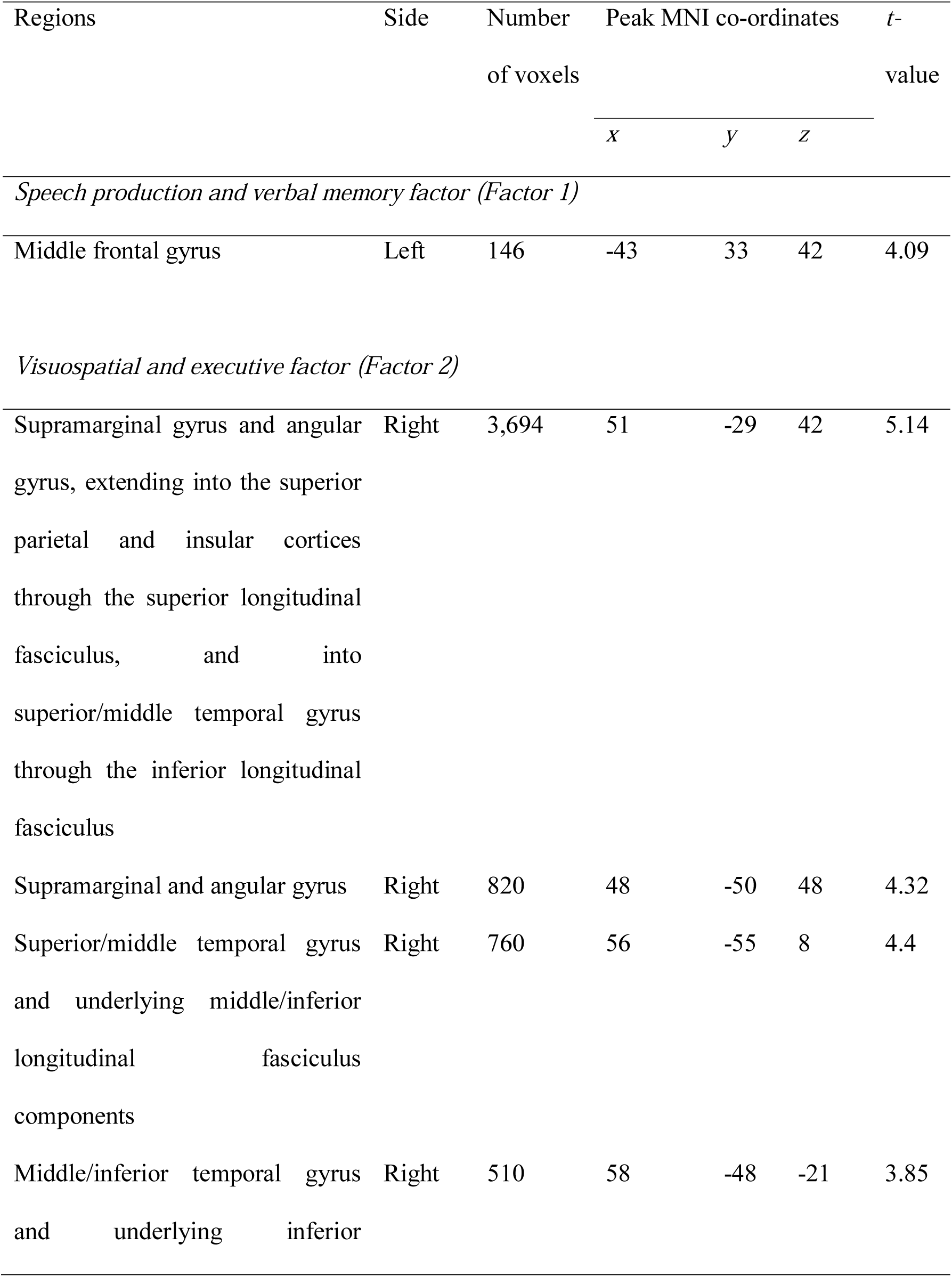
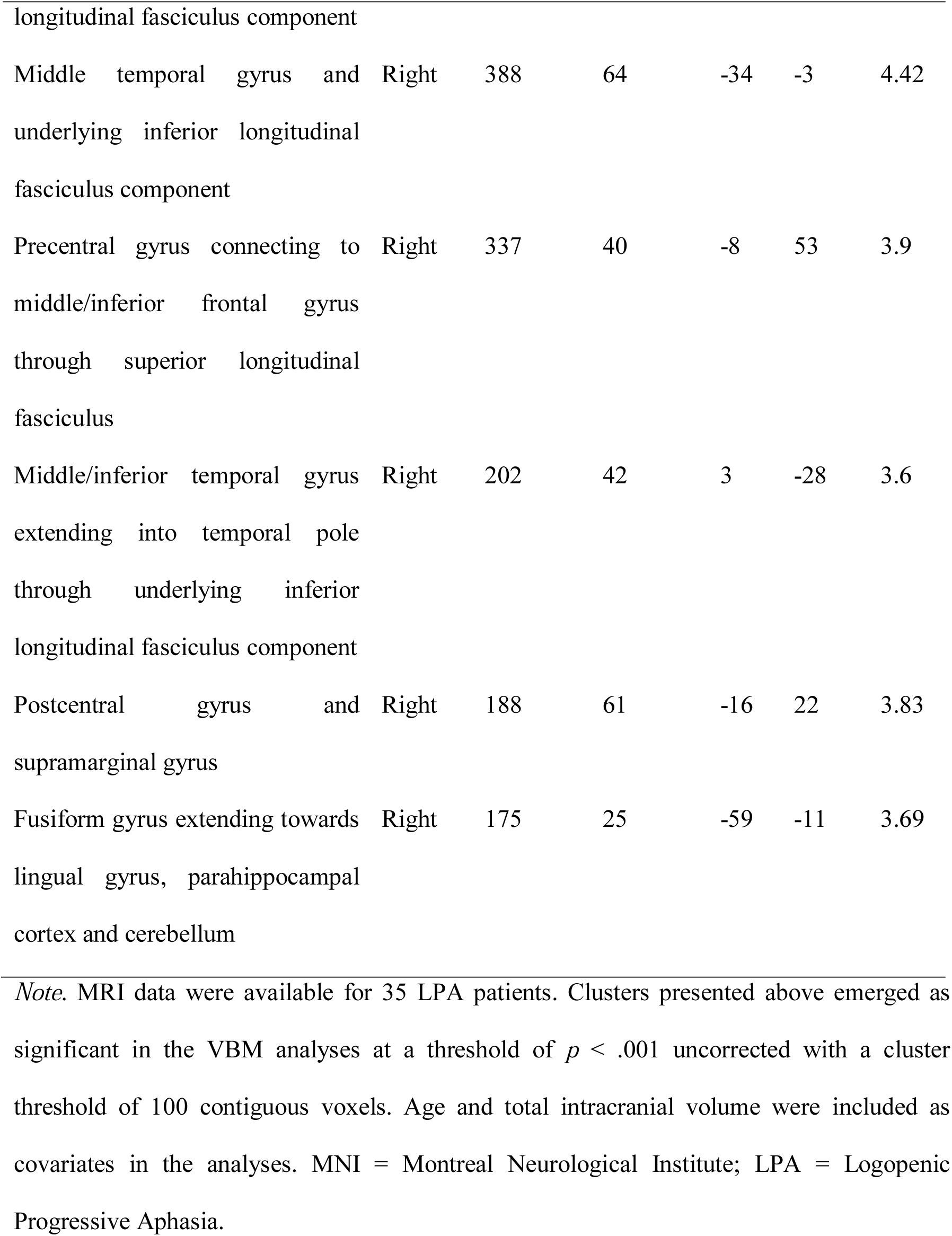
Voxel-based morphometry results showing regions of grey and white matter intensity that correlate with PCA-generated Factor 1 and Factor 2 scores in the LPA group.

**Fig 5.**
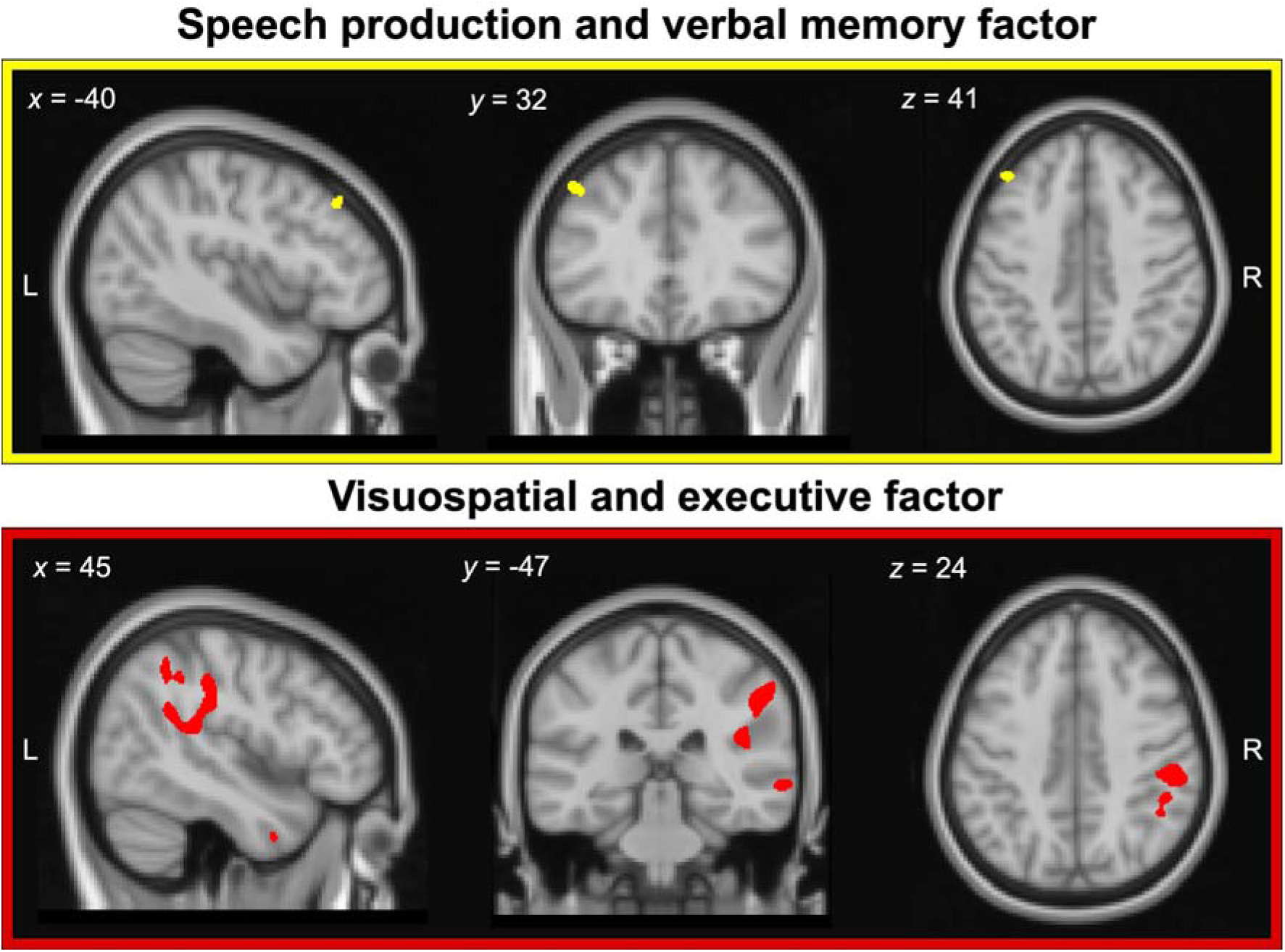
Regions of grey and white matter intensity that uniquely correlate with factor scores on the speech production and verbal memory factor (*i.e*., Factor 1; upper panel) and visuospatial and executive factor (*i.e*., Factor 2; lower panel) in LPA patients. Both factors were derived from varimax-rotated PCA of neuropsychological test performance in the LPA group. Coloured voxels indicate regions that emerged significant in the voxel-based morphometry analyses at a threshold of *p* < .001 uncorrected for multiple comparisons with a cluster threshold of 100 contiguous voxels. All clusters reported at *t* = 4.09 for speech production and verbal memory factor and *t* ≥ 3.6 for visuospatial and executive factor. Age and total intracranial volume were included as covariates in the analyses. Clusters are overlaid on the Montreal Neurological Institute (MNI) standard brain with *x, y*, and *z* co-ordinates reported in MNI standard space. L = Left; R = Right; PCA = Principal Component Analysis; LPA = Logopenic Progressive Aphasia.

##### 3.6.4.1. Speech production and verbal memory factor (Factor 1)

In the overall LPA group, speech production and verbal memory factor scores were found to correlate with grey matter intensity of the left middle frontal gyrus (Table 3, Figure 5, upper panel).

##### 3.6.4.2. Visuospatial and executive factor (Factor 2)

Visuospatial and executive factor scores in LPA correlated with grey and white matter intensity in right lateral parietal (supramarginal gyrus, angular gyrus) and medial parietal (precentral and postcentral gyri), right lateral temporal regions (superior/middle/inferior temporal gyri) and the right middle frontal gyrus. Additionally, a small cluster in the ventral temporal cortex (fusiform, lingual, and parahippocampal gyrus) extending into the right cerebellar cortex was noted. Changes in white matter intensity of the right superior longitudinal fasciculus (connecting frontoparietal cortices) and right middle/inferior longitudinal fasciculus (connecting temporoparietal cortices) were further found to correlate with visuospatial and executive factor scores (Table 3, Figure 5, lower panel).

In summary, both factors were found to correlate with distinct neural regions, with the speech production and verbal memory factor scores (Factor 1) correlating with grey matter intensity of the middle frontal gyrus, and the visuospatial and executive factor scores (Factor 2) correlating with largely right-sided temporoparietal and frontal regions and their underlying white matter connections. Importantly, the regions to emerge as significant in our covariate analyses (Figure 5) are not the areas of maximal atrophy in LPA (Figure 4A) but rather those with greater variance in grey and white matter intensity (Figure 4B and 4C) which flank the areas of maximal atrophy.

## 4. DISCUSSION

This study demonstrates that the presence of visuospatial and executive deficits in LPA, beyond core language disturbance, does not reflect advancing disease severity. Instead, these deficits in LPA form their own independent cognitive dimension with discrete neuroanatomical bases and are reliably present even in the early stages of LPA. In more detail, the PCA identified two emergent factors capturing the heterogeneity of the LPA cognitive profile. The first factor reflected expressive language and phonological working memory impairments that are not only diagnostic of LPA (Gorno-Tempini et al., 2008) but hold discriminative ability in differentiating LPA from other primary progressive aphasia syndromes (Gorno-Tempini et al., 2011). Our findings mesh well with previous studies employing other data-driven approaches such as two-step and hierarchical clustering analyses in LPA (Leyton et al., 2015; Machulda et al., 2013; Owens et al., 2018) and confirm that verbal working memory, repetition and naming difficulties typify this syndrome.

Importantly, however, our PCA approach revealed a second, orthogonal factor comprising nonverbal episodic memory, visuo-constructional, attentional and executive processing, as well as receptive language and comprehension measures. This visuospatial and executive factor was independent of expressive language difficulties in LPA, running counter to the view that ‘general cognitive’ impairment in LPA reflects little more than the language demands of neuropsychological measures (Machulda et al., 2013; Owens et al., 2018). In addition, we found that performance deficits on this second factor were pervasive across the entire LPA cohort, regardless of the severity of their language impairments. Again, this finding is not easily accommodated by previous proposals that global cognitive decline in LPA is a product of advancing disease severity (Funayama et al., 2013; Machulda et al., 2013; Owens et al., 2018). Rather, our findings indicate the presence of a genuine co-occurring global cognitive impairment, spanning multiple domains, that is independent of language function and disease severity. This view is in keeping with recent findings of marked nonverbal memory and emotion processing disturbances, even after accounting for expressive language impairments and disease severity in LPA (Multani et al., 2017; Ramanan et al., 2016). More generally, these results add to the view that subtypes of Alzheimer’s disease reflect graded rather than absolute variations presumably reflecting individual differences in the exact distribution of Alzheimer’s pathology (c.f., Lambon Ralph et al., 2003).

At an individual-level, systematic variations on the visuospatial and executive factor, regardless of patient performance on the language factor, underline at the graded nature of the changes across patients. Adopting a case-comparison approach, we demonstrated that two LPA patients with comparable expressive language impairment (determined on Factor 1) diverge considerably on their visuo-executive performance. Importantly, this pattern was present even when comparing pairs of LPA patients with mild, moderate, or severe language difficulties, suggesting attention, executive and visuospatial deficits are core features of the LPA syndrome. From a clinical standpoint, our findings align well with previous descriptions of single cases of LPA presenting with ‘atypical’ symptoms. For example, single cases of LPA have been described to present with a marked breakdown in attentional processing manifesting in hemi-spatial neglect (Zilli & Heilman, 2016). Similarly, individuals with LPA have been described as presenting with profound and co-occurring visuospatial disturbances notable in judging distances and reach-to-grasp difficulties (Fitzpatrick, Blanco-Campal, & Kyne, 2019). Importantly, these ‘atypical’ symptoms emerged in the context of otherwise language deficits and atrophy profiles typical of LPA (Fitzpatrick et al., 2019; Zilli & Heilman, 2016). Our case-comparison findings indicate that marked individual-level variability in non-linguistic cognitive performance is a key feature of LPA and suggest caution in excluding cases who present with such early co-occurring deficits.

We next explored associations between factor scores and cortical and subcortical brain changes in LPA. Performance on the speech production and verbal memory factor was found to correlate with grey and white matter changes of the left middle frontal gyrus. This region is a key frontal node of the language and executive processing networks, with well described roles in supporting fluency in expressive language (Abrahams et al., 2003; Rogalski, Cobia, Harrison, Wieneke, Thompson, et al., 2011) and working memory (Whitwell, Jones, et al., 2015). In particular, middle frontal, along with neighbouring prefrontal cortical regions are posited to play a role in maintaining information within working memory (D’Esposito & Postle, 1999). Disrupted functional connectivity of the middle frontal gyrus with prefrontal, lateral and medial parietal regions has been linked to working memory impairments in LPA (Whitwell, Jones, et al., 2015), with cortical thickness of this region further associated with reduced verbal fluency (as measured by mean length of utterance during story telling) in patients with primary progressive aphasia (Rogalski, Cobia, Harrison, Wieneke, Thompson, et al., 2011). Although not typical of the early LPA atrophy pattern, middle frontal gyrus atrophy has been described previously in the syndrome (Phillips et al., 2019; Rohrer et al., 2010), and tends to become more salient as atrophy progresses along the left sylvian fissure into fronto-insular regions (Rohrer et al., 2013). It is possible, therefore, that this middle frontal region shows greater inter-participant variance and thus greater sensitivity to detect associations in the VBM correlation analyses. This is in contrast to the left temporoparietal cortices which are atrophied early and consistently in LPA patients, and thus, resultantly, have low atrophy variance across the group. Future explorations of the temporal unfolding of cortical atrophy patterns and their inter-participant variance, in relation to the cognitive profiles outlined here will be important.

Turning our attention to Factor 2, performance on the visuospatial and executive factor was found to correlate with grey and white matter intensity of right-lateralised temporoparietal and prefrontal regions, including precentral, inferior parietal, lateral temporal, inferior frontal and insular cortices. Moreover, LPA patients with poorer scores on the visuospatial and executive factor tended to demonstrate greater right-lateralised temporoparietal and prefrontal atrophy. Right-lateralised regions such as precentral gyrus and superior/inferior parietal regions are typically proposed to regulate goal-directed and stimulus-driven attentional abilities (Corbetta & Shulman, 2002), while middle/inferior frontal regions have been noted to aid in executive processing by regulating control and inhibitory functions (Aron, Robbins, & Poldrack, 2004; Sridharan, Levitin, & Menon, 2008), respectively. Frontotemporal dementia patients with increased pathological burden to the right hemisphere tend to show greater impairment on nonverbal measures of executive functioning, when compared to those with left-lateralised atrophy (Boone et al., 1999). A similar pattern has been noted in LPA, wherein the presence of right-hemisphere frontal and temporoparietal atrophy reliably signals the emergence of attentional, executive, and general cognitive impairments in the syndrome (Machulda et al., 2013). Similarly, although impairment in single-word comprehension currently forms an exclusion criterion for the diagnosis of LPA (Gorno-Tempini et al., 2011), recent studies incorporating *in vivo* confirmation of underlying Alzheimer’s pathology revealed marked single word comprehension difficulties in LPA (Leyton et al., 2015; Louwersheimer et al., 2016). In fact, LPA patients with single word comprehension impairment tend to demonstrate greater atrophy to right-lateralised temporal regions, centred on the fusiform, and inferior/middle temporal cortices (Faria, Sebastian, Newhart, Mori, & Hillis, 2014; Leyton et al., 2015). We speculate that encroachment of atrophy into right temporoparietal and prefrontal grey/white matter may predict the onset of visuospatial and executive performance impairments in LPA, however, longitudinal studies will be crucial to test this proposal.

The current findings must be interpreted in the context of certain caveats. First, the majority of our LPA patients have not yet come to autopsy, precluding confirmation of underlying Alzheimer versus non-Alzheimer pathology in our cohort. Nevertheless, we rigorously applied the diagnostic criteria of LPA (Gorno-Tempini et al., 2011) to ensure the exclusion of other primary progressive aphasia syndromes presenting with primary semantic processing or grammatical impairments. Studies employing PCA approaches necessarily rely upon the nature of data fed into the model. Given that this was a retrospective study, we were constrained by the cognitive measures available to us, however, we included detailed standardised measures of multiple cognitive domains, leading to findings that, in the context of the existing literature, make intuitive sense. Given emerging evidence of behavioural changes in LPA (e.g., increased reports of anxiety; Magnin et al., 2013), future studies will benefit from exploring if behavioural changes in LPA occur independently of language impairment in the syndrome or co-occur with the visuospatial and executive factor identified here. Of further importance is the need to establish associations between cognitive factors and underlying pathological markers in LPA, given extant evidence for distinct patterns of cognitive performance and lateralized deposition of underlying pathology in LPA patients with underlying Alzheimer versus non-Alzheimer pathology (Giannini et al., 2017; M. Mesulam et al., 2008; Whitwell, Duffy, et al., 2015). Finally, we reported our VBM results at an uncorrected threshold of *p* < .001, however, this threshold is far more conservative than traditional multiple comparison approaches such as false discovery rate, and is increasingly used when exploring links between cognition and neurodegeneration (Sheelakumari et al., 2019; Whitwell et al., 2010).

Despite these limitations, our findings hold important clinical implications relevant to the diagnosis and characterisation of LPA. Identification of heterogeneity in cognitive function in LPA underscores the need for comprehensive neuropsychological workup beyond language in primary progressive aphasia. By limiting their primary focus to language impairments, clinicians will underestimate the presence and severity of visuospatial and executive impairments in LPA, potentially leading to increased functional disturbances and carer burden. We further speculate that the emergence of visuospatial and executive impairments in LPA can be thought of as converse to atypical variants of Alzheimer’s disease such as Posterior Cortical Atrophy. Although described as a syndrome with preponderant visual disturbances due to early right-sided parietal atrophy, Posterior Cortical Atrophy patients gradually demonstrate increasing language and verbal working memory dysfunction (Crutch, Lehmann, Warren, & Rohrer, 2013; Trotta, Lamoureux, Bartolomeo, & Migliaccio, 2019). This would suggest the existence of a possible continuum between these syndromes, with LPA unfolding to resemble Posterior Cortical Atrophy later in the disease course (Fitzpatrick et al., 2019). More generally, these collective results might imply that typical Alzheimer’s disease and its multiple atypical subtypes might all be reconceptualised in terms of graded variations within a single multiple dimensional space (Lambon Ralph et al., 2003). Future studies directly comparing the cognitive, behavioural and neural trajectories of these syndromes over time will be critical to address this question.

In conclusion, we provide new insights into the syndrome of LPA, by revealing a fundamental impairment of visuospatial and executive processes, independent of the characteristic language difficulties in this syndrome. This visuospatial and executive impairment varies systematically across LPA patients, irrespective of disease severity, and correlates with right-lateralised temporoparietal and frontal regions. Our findings reveal the inherent complexity of the LPA syndrome in terms of cognitive profiles and neural atrophy patterns and suggest that reconceptualization of the LPA syndrome and its relationship to typical and atypical variants of Alzheimer’s disease is warranted.

## 5. Acknowledgements

The authors are grateful to the patients and families for their continued support of our research. The authors wish to acknowledge the Sydney Informatics Hub funded by The University of Sydney for providing access to High Performance Computing (HPC) facilities.

## Supplementary material

One supplementary material with supplementary methods, four supplementary tables and five supplementary figures

