## Supplementary for "Establishing two principal dimensions of cognitive variation in Logopenic Progressive Aphasia"

**SUPPLEMENTARY METHODS**

**Statistical analyses**

**Converting raw scores to percentages**

Where tests followed a linear scoring pattern (with higher scores indicative of better performance) and had maximum scores attainable (e.g., ACE-R, Digit Span, SYDBAT, and ROCF measures), data were converted into percentages as follows: [(raw score / max score) * 100]. Tests like the TMT B-A, on the other hand, followed an inverse scoring pattern (with higher scores indicative of poorer performance) with no maximum attainable scores; a two-step approach to standardization was hence adopted. First, TMT B-A scores were converted into percentages using the ‘local maximum’ score (*i.e.*, highest TMT B-A score, indicative of poorest performance) using the formula: (raw score / local maximum) * 100. This score was subsequently subtracted from 100 to derive an index of TMT B-A performance percentage.

**Imputing missing data using Probabilistic Principal Components Analysis**

Missing data were imputed using a probabilistic PCA approach. Briefly, this approach estimates principal components in the entire dataset (including missing data) and uses this information to predict missing values in a probabilistic manner. As the end result produces a ‘full’ dataset with no missing values, probabilistic PCA approaches offer improved stability as compared to list-wise exclusion of rows with missing data, while simultaneously guarding against overfitting of imputed data points (unlike imputation of group mean) (see Tipping and Bishop, 1999; Ilin and Raiko, 2010).

Here, we used a *k*-fold cross-validation approach (with *k* = 4) to impute missing data. In this approach, the dataset was first divided into three ‘training’ subsets and one ‘testing’ subset (as *k* = 4). Following this, a *k*-fold cross-validation approach with 1,000 permutations was used to choose the component solution (i.e., number of components) that optimally represent the underlying data structure. The most ‘stable’ solution with the lowest root-mean-squared-error for all held-out cases (i.e., cases in the testing set) over 1,000 permutations was used (Ballabio, 2015). This approach was also used to select the optimal number of components for subsequent PCA using the imputed dataset. All imputed scores were visually inspected by S.R. to ensure they fit with each patient’s overall cognitive profile.

**Projecting lower bound of normality from Control data into the patient PCA-space**

First, all Control data were standardized into percentages. For each test, the ‘lower bound of normality’ (-1.96 standard error of the mean) was estimated, and *z*-scored using the mean and standard deviation of the respective test. Then, the *z*-scored lower bound of normality value was multiplied with its respective component coefficient (*i.e.*, loading) derived from the PCA run in the LPA group to receive a product. This product reflected the Control lower bound of normality for each measure included in the patient PCA. As we were interested in a single value representative of the lower bound of normality for each of the PCA-generated factors, we retained only products for tests loading highly (loadings > .5) on each factor. The retained products were summed to derive a single value reflecting the Control lower bound of normality in the patient PCA space for that specific factor.

**VBM analyses**

**Pre-processing**

A standardized pre-processing pipeline was employed wherein images were first segmented into three tissue probability maps (grey matter, white matter, cerebrospinal fluid) using the segmentation option implemented in SPM12. Grey and white matter probability maps were then normalised with the in-built Diffeomorphic Anatomical Registration using Exponentiated Lie Algebra (DARTEL: Ashburner, 2007) approach. A study-specific template was then computed using all the grey and white matter probability maps, following which, these maps were spatially normalised to Montreal Neurological Institute (MNI) space using the transformation parameters from the corresponding DARTEL template. Finally, images were modulated and smoothed with an 8mm full-width-half-maximum Gaussian kernel to increase signal-to-noise ratio. The segmented, normalised, modulated and smoothed grey and white matter images for all participants were finally concatenated into a 4-dimensional combined grey-white matter image for subsequent VBM analyses. To determine anatomical labels, the Johns Hopkins University White Matter atlas and the ICBM-DTI-WM atlas labels were used (Mori *et al.*, 2008; Oishi *et al.*, 2008).

**Supplementary Table 1**. Correlations (*r* values with exact *p* values indicated in parenthesis) between neuropsychological test performance and a clinician-indexed disease severity score in the LPA group

| Tests | CDR-FTLD SoB |
| --- | --- |
| ACE-R Attention | -.29 (.059) |
| ACE-R Memory | -.28 (.06) |
| ACE-R Fluency | -.18 (.251) |
| ACE-R Language | -.02 (.85) |
| ACE-R Visuospatial | -.23 (.140) |
| SYDBAT Naming | -.07 (.632) |
| SYDBAT Comprehension | .02 (.855) |
| SYDBAT Repetition | -.24 (.119) |
| SYDBAT Semantic Association | .02 (.880) |
| **Digit Span Forward** | **-.39 (.010)** |
| Digit Span Backward | -.15 (.319) |
| ROCF Copy | -.21 (.196) |
| ROCF Delayed Recall | -.06 (.686) |
| TMT B-A | .04 (.844) |

*Note*. All correlations computed using Pearson’s method corrected for multiple comparisons using ‘false discovery rate’ approach; ACE-R = Addenbrooke’s Cognitive Examination – Revised; SYDBAT = Sydney Language Battery; ROCF = Rey-Osterrieth Complex Figure; TMT B-A = Trail Making Test parts B-A; LPA = Logopenic Progressive Aphasia.

**Supplementary Table 2**. Factor scores for all LPA patients on varimax-rotated Principal Components Analysis of neuropsychological data

| Patient number | Speech production and verbal memory factor (Factor 1) | Visuospatial and executive factor (Factor 2) |
| --- | --- | --- |
| 1 | -1.146 | -1.64511 |
| 2 | -0.24616 | 1.570906 |
| 3 | -2.24227 | -1.01342 |
| 4 | 0.485719 | -0.12502 |
| 5 | 1.098248 | 0.297067 |
| 6 | -0.74277 | 1.041495 |
| 7 | -0.89471 | 0.880863 |
| 8 | -0.2461 | -1.68694 |
| 9 | 0.688448 | -0.68057 |
| 10 | -0.97227 | -1.11695 |
| 11 | -0.76827 | 0.180948 |
| 12 | 0.186963 | -0.69045 |
| 13 | 0.774586 | 0.330748 |
| 14 | -1.37612 | -0.86472 |
| 15 | -0.61102 | -0.20566 |
| 16 | 0.533649 | -0.53612 |
| 17 | 0.169346 | 0.977779 |
| 18 | 0.392762 | 0.715236 |
| 19 | 1.726422 | 0.237475 |
| 20 | 0.647877 | 0.510045 |
| 21 | 2.036857 | 0.17811 |
| 22 | -0.43836 | 0.835594 |
| 23 | -0.16357 | 0.53522 |
| 24 | -1.76387 | 0.436377 |
| 25 | 0.268567 | 0.465436 |
| 26 | -0.53805 | 0.765265 |
| 27 | 0.376161 | -0.8835 |
| 28 | -1.42891 | -0.84649 |
| 29 | 0.539608 | -1.86 |
| 30 | -0.73812 | 1.028097 |
| 31 | -0.74169 | 2.639473 |
| 32 | 1.0412 | 0.833565 |
| 33 | 0.173351 | -1.48031 |
| 34 | -1.25718 | 1.911062 |
| 35 | 1.331319 | -0.16142 |
| 36 | -0.04695 | 0.316125 |
| 37 | 1.308572 | -0.44626 |
| 38 | 0.473009 | 0.175668 |
| 39 | 1.088622 | -0.40421 |
| 40 | 1.994399 | -0.28698 |
| 41 | 0.021618 | -0.4279 |
| 42 | -0.99269 | -1.83626 |
| 43 | -0.00221 | 0.335747 |

*Note*. Measures loading heavily (loadings > .5) on the speech production and verbal memory factor (Factor 1) comprised the ACE-R Language and Memory subdomains, SYDBAT Repetition and Naming subtests, and Digit Span Forward and Backward tests. Measures loading heavily on the visuospatial and executive factor (Factor 2) comprised the ACE-R Attention subtest, SYDBAT Semantic and Comprehension subtests, ROCF Copy and Delayed Recall measures, and TMT B-A measure. LPA = Logopenic Progressive Aphasia; ACE-R = Addenbrooke’s Cognitive Examination Revised; SYDBAT = Sydney Language Battery; ROCF = Rey-Osterrieth Complex Figure; TMT B-A = Trail Making Test parts B – A.

**Supplementary Table 3**. Voxel-based morphometry results showing regions of significant grey and white matter intensity decrease in the LPA group relative to Controls

| Contrast | Regions | Side | Number of voxels | Peak MNI co-ordinates | | | *t*-value |
| --- | --- | --- | --- | --- | --- | --- | --- |
|  |  |  |  | *x* | *y* | *z* |  |
| LPA vs. Controls | Bilateral superior/middle/inferior temporal gyrus extending towards left supramarginal gyrus and left angular gyrus through the left inferior longitudinal/inferior fronto-occipital fasciculus, bilateral precuneus and bilateral middle/posterior cingulate cortex connecting to bilateral hippocampus, bilateral parahippocampal gyrus through the cingulate bundle, bilateral thalamus including anterior thalamic radiation, bilateral amygdala, bilateral temporal pole into right orbitofrontal cortex through uncinate fasciculus, bilateral insular cortex, and left superior/middle frontal gyrus and underlying superior longitudinal fasciculus component | B | 297,437 | -48 | -45 | -21 | 11.17 |
|  | Superior/middle frontal gyrus and underlying superior longitudinal fasciculus component | R | 3,292 | 26 | 56 | 8 | 7.32 |
|  | Anterior/middle cingulate cortex connecting to superior/middle frontal gyrus through superior longitudinal fasciculus and cingulum (cingulate gyrus) bundles | R | 3,008 | 24 | 29 | 36 | 6.87 |
|  | Right superior/middle temporal gyrus extending towards angular gyrus and lateral occipital cortex involving inferior longitudinal fasciculus | R | 1,069 | 55 | -62 | 12 | 8.15 |
|  | Fusiform gyrus, middle/inferior occipital gyrus, lingual gyrus, cerebellum | L | 411 | -31 | -77 | -16 | 5.87 |
|  | Superior/middle frontal gyrus and underlying superior longitudinal fasciculus component | L | 227 | -2 | 49 | 35 | 6.11 |

*Note.* MRI data for VBM analyses were available for 63 participants (35 LPA, 28 Controls). Clusters presented above emerged as significant in the VBM analyses at *p* < .01 corrected for Family-Wise Error with a cluster threshold of 100 contiguous voxels. Age and total intracranial volume were included as covariates in the analyses. MNI = Montreal Neurological Institute; LPA = Logopenic Progressive Aphasia; L = Left; R = Right; B = Bilateral.

**Supplementary Table 4**. Voxel-based morphometry results showing regions of significant grey and white matter intensity decrease in LPA patients split on Factor 1 and Factor 2 scores

| Contrast^†^ | Regions | Side | Number of voxels | MNI coordinates | | | *t*-value |
| --- | --- | --- | --- | --- | --- | --- | --- |
|  |  |  |  | *x* | *y* | *z* |  |
| **Split on Factor 1** | | | | | | | |
| Positive > Negative | No significant clusters |  |  |  |  |  |  |
| Positive > Negative | No significant clusters |  |  |  |  |  |  |
| **Split on Factor 2** | | | | | | | |
| Positive > Negative | No significant clusters |  |  |  |  |  |  |
| Negative > Positive | Angular gyrus, supramarginal gyrus, superior/middle temporal gyrus extending into insular cortex through underlying superior longitudinal fasciculus | R | 8,713 | 49 | -50 | 48 | 5.33 |
|  | Superior/middle temporal gyrus and underlying middle/inferior longitudinal fasciculus bundle | R | 1,347 | 59 | -56 | 8 | 5.5 |
|  | Posterior cingulate cortex and precuneus | R | 923 | 17 | -53 | 12 | 4.62 |
|  | Occipital cortex extending into cerebellum | R | 779 | 32 | -52 | -16 | 3.91 |
|  | Angular gyrus and underlying superior/inferior longitudinal fasciculus component | L | 596 | -46 | -61 | 46 | 4.53 |
|  | Middle frontal gyrus and underlying superior longitudinal fasciculus component | R | 574 | 46 | 34 | 28 | 3.92 |
|  | Middle cingulate cortex | R | 527 | 2 | -19 | 36 | 4.05 |
|  | Middle frontal gyrus and underlying superior longitudinal fasciculus component | R | 337 | 38 | -4 | 51 | 3.69 |
|  | Precuneus, posterior/middle cingulate gyrus and underlying cingulum (cingulate gyrus) bundle | R | 329 | 1 | -46 | 45 | 4.68 |
|  | Middle/inferior frontal gyrus and underlying middle/inferior longitudinal fasciculus component | R | 296 | 50 | 40 | 9 | 4.51 |
|  | Precuneus | L | 252 | -4 | -50 | 56 | 3.96 |
|  | Superior/middle temporal gyrus and inferior longitudinal fasciculus | R | 232 | 24 | 50 | 36 | 3.96 |
|  | Superior/middle/inferior temporal gyrus and underlying inferior longitudinal fasciculus component | R | 188 | 44 | -1 | -49 | 3.56 |
|  | Angular gyrus, superior/middle temporal gyrus | R | 162 | 52 | -67 | 29 | 3.77 |
|  | Fusiform gyrus, parahippocampal gyrus, inferior temporal gyrus | R | 156 | 41 | -24 | -21 | 3.72 |
|  | Middle frontal gyrus and underlying superior longitudinal fasciculus component | R | 148 | 24 | 35 | -13 | 3.6 |

*Note.* MRI data available for 35 LPA patients. ^†^For each factor, positive indicates patients with higher factor scores (*i.e.*, greater than zero) and negative indicates patients with lower factor scores (*i.e.*, less than zero). Clusters presented emerged as significant in the VBM analyses *p* < .001 uncorrected with a cluster threshold of 100 contiguous voxels. Age and total intracranial volume were included as covariates in the analyses. MNI = Montreal Neurological Institute; LPA = Logopenic Progressive Aphasia; L = Left; R = Right.

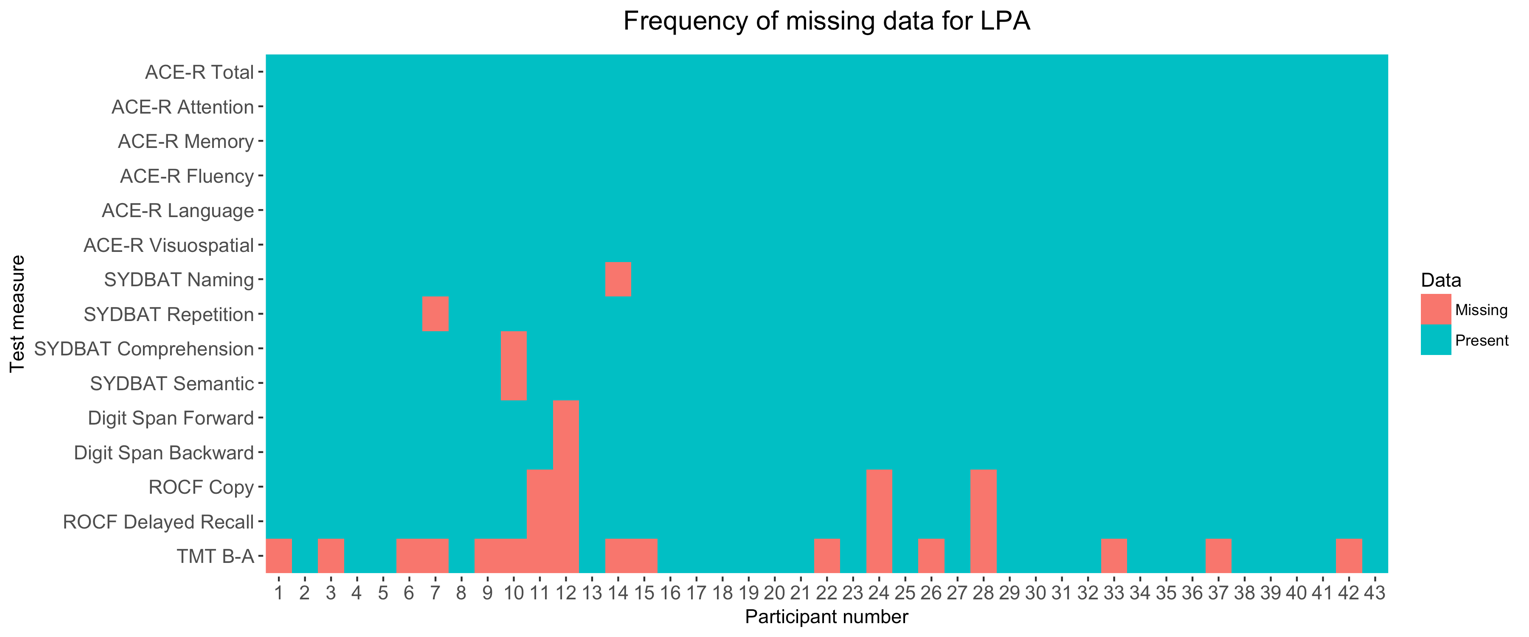

**Supplementary Figure 1**. Frequency of missing data (overall 4.8%, indicated in orange) on neuropsychological assessments for LPA patients. LPA = Logopenic Progressive Aphasia; ACE-R = Addenbrooke’s Cognitive Examination – Revised; SYDBAT = Sydney Language Battery; ROCF = Rey-Osterrieth Complex Figure; TMT B-A = Trail Making Test parts B-A.

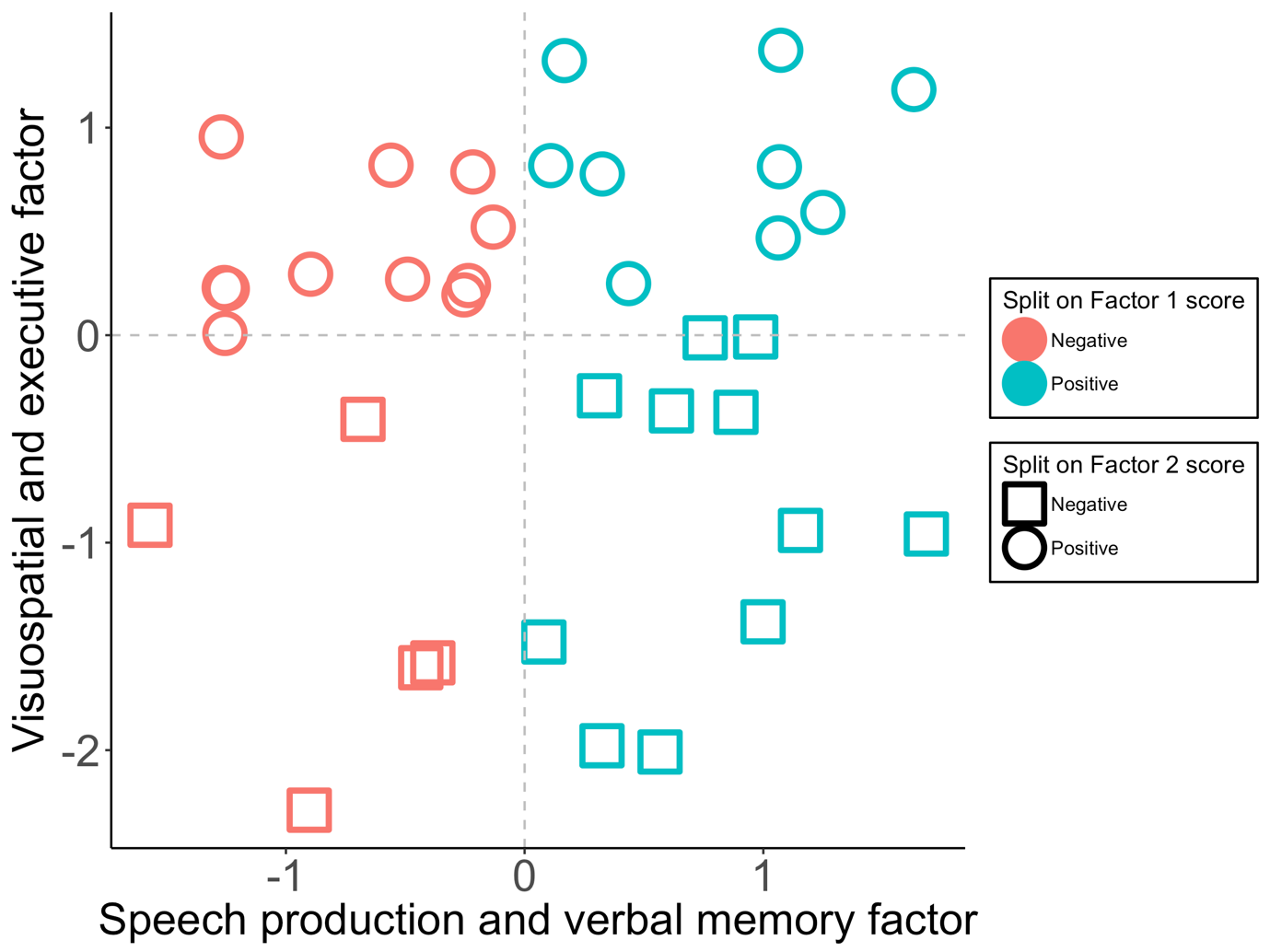

**Supplementary Figure 2**. The LPA group split as per their scores on positive or negative side of a zero score on Factors 1 and 2. Only LPA patients with MRI data (*N* = 35) are displayed. Measures loading heavily (loadings > .5) on the speech production and verbal memory factor (Factor 1) included the ACE-R Language and Memory subdomains, SYDBAT Repetition and Naming subtests, and Digit Span Forward and Backward tests. Measures loading heavily on the visuospatial and executive factor (Factor 2) included the ACE-R Attention subtest, ROCF Copy and Delayed Recall measures, TMT B-A measure, and SYDBAT Semantic and Comprehension subtests. LPA = Logopenic Progressive Aphasia; ACE-R = Addenbrooke’s Cognitive Examination Revised; SYDBAT = Sydney Language Battery; ROCF = Rey-Osterrieth Complex Figure; TMT B-A = Trail Making Test parts B – A.

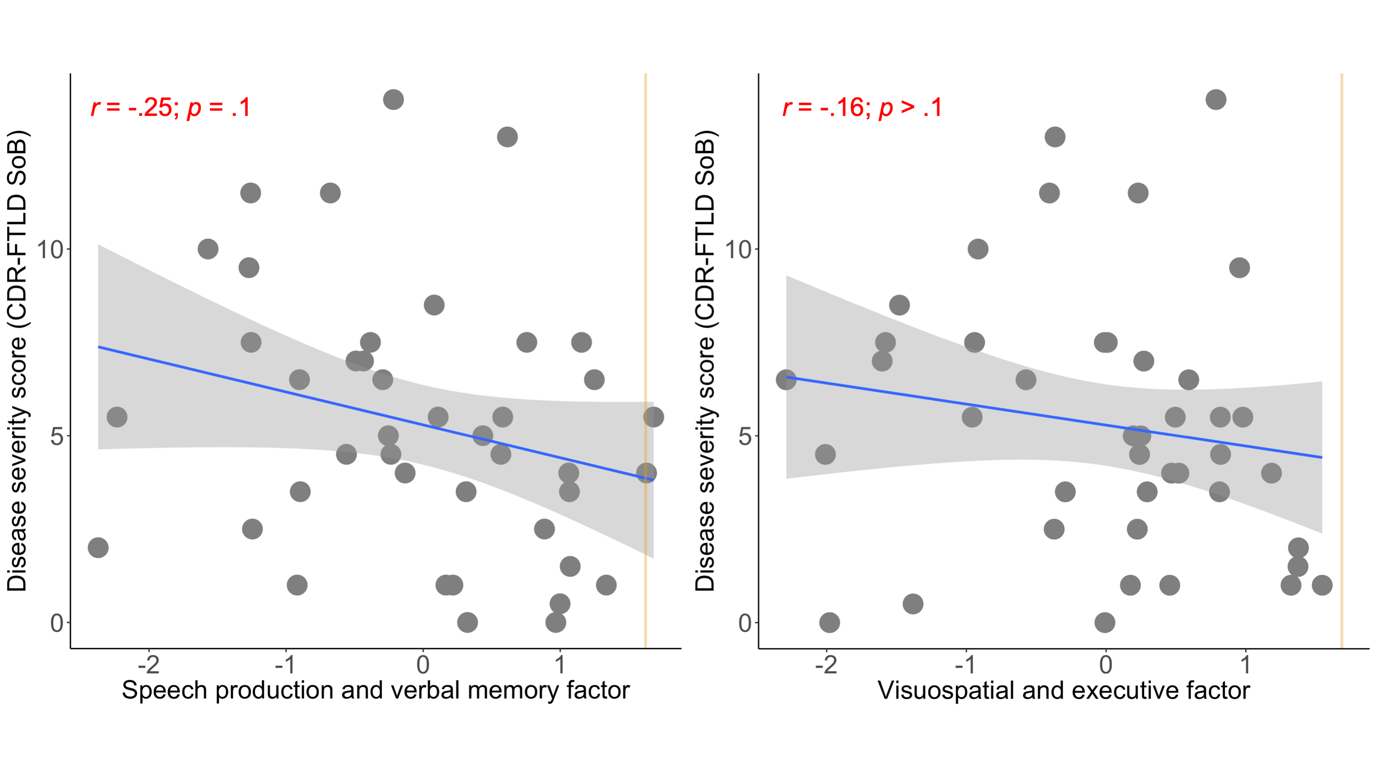

**Supplementary Figure 3.** Pearson’s correlations between disease severity (measured using CDR-FTLD SoB score) and emergent factor scores from PCA for Factors 1 (left panel) and Factor 2 (right panel) within the LPA group. Grey bands indicate standard error. Gold lines indicate lower bound of normality (-1.96 standard error from the mean) as estimated from the Control group (calculation detailed in Supplementary Methods). CDR-FTLD SoB = Clinical Dementia Rating – Frontotemporal Lobar Degeneration Sum of Boxes; LPA = Logopenic Progressive Aphasia.

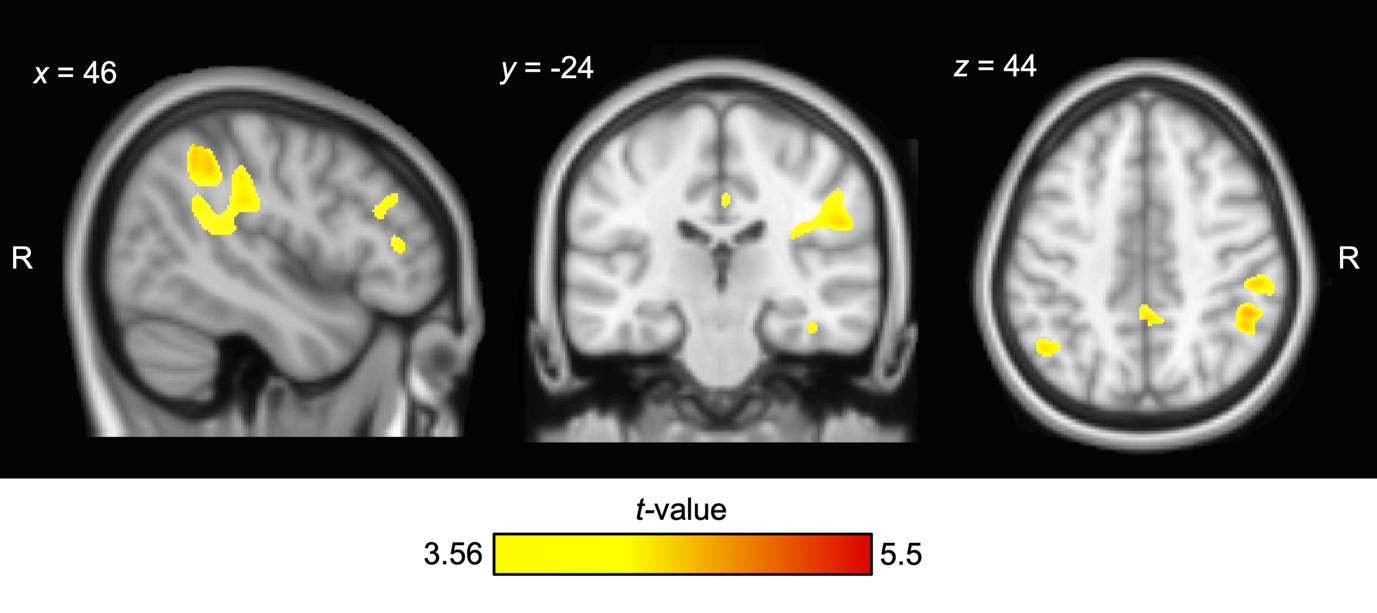
**Supplementary Figure 4**. Voxel-based morphometry analyses of combined grey and white matter showing regions of significant grey and white matter intensity reduction in LPA patients with negative Factor 2 scores (*i.e.*, worse performance on Factor 2) as compared to LPA patients with positive Factor 2 scores (*i.e.*, better performance on Factor 2). Coloured voxels indicate regions that emerged significant in the voxel-based morphometry analyses at *p* < .001 uncorrected with a cluster threshold of 100 contiguous voxels. Age and total intracranial volume were included as covariates in the analyses. Clusters are overlaid on the Montreal Neurological Institute (MNI) standard brain with *x*, *y*, and *z* co-ordinates reported in MNI standard space. R = Right; LPA = Logopenic Progressive Aphasia.
